# *BCOR* and *BCORL1* mutations disrupt PRC1.1 repressive function in leukemia by unlinking the RING-PCGF1 enzymatic core from target genes

**DOI:** 10.1101/2021.03.08.433705

**Authors:** Eva J. Schaefer, Helen C. Wang, Clifford A. Meyer, Paloma Cejas, Micah D. Gearhart, Emmalee R. Adelman, Iman Fares, Annie Apffel, Klothilda Lim, Yingtian Xie, Christopher J. Gibson, Monica Schenone, H. Moses Murdock, Eunice S. Wang, Lukasz P. Gondek, Martin P. Carroll, Rahul S. Vedula, Eric S. Winer, Jacqueline S. Garcia, Richard M. Stone, Marlise R. Luskin, Steven A. Carr, Henry W. Long, Vivian J. Bardwell, Maria E. Figueroa, R. Coleman Lindsley

## Abstract

*BCOR* and its paralog *BCORL1* encode subunits of the Polycomb repressive complex 1.1 (PRC1.1) and are recurrently mutated in myeloid malignancies. We show that leukemia-associated *BCOR/BCORL1* mutations unlink the PRC1.1 RING-PCGF enzymatic core from the KDM2B-containing chromatin targeting auxiliary subcomplex, either by causing complete protein loss or expression of a C-terminally truncated protein lacking the PCGF Ub-like fold discriminator (PUFD) domain. By uncoupling PRC1.1 repressive function from target genes, *BCOR/BCORL1* mutations activate aberrant cell signaling programs that confer acquired resistance to treatment. This study provides a mechanistic basis for Polycomb repressive dysfunction as a key oncogenic driver in myeloid malignancies and identifies a potential strategy for targeted therapy in *BCOR*-mutated cancer.

## Introduction

Polycomb repressive complex 1 (PRC1) is a modularly assembled multisubunit complex that controls programmatic gene expression via effects on histone epigenetic marks. PRC1 is defined by having a RING-PCGF enzymatic core, whose main activity is to monoubiquitinate H2A on lysine 119 (Blackledge et al., 2020; Gao et al., 2012; Tamburri et al., 2020). PRC1 can assemble as 6 different complexes (PRC1.1-PRC1.6), each defined by a specific PCGF linker protein that couples RING1/RNF2 enzymes to a distinct set of auxiliary subunits that determine its mechanism of chromatin targeting and gene regulatory function (Gao et al., 2012; Scelfo et al., 2019). Of these six PRC1 subcomplexes, only PRC1.1 is known to be somatically mutated in cancer (Sashida et al., 2019).

In PRC1.1, the enzymatic core is linked by PCGF1 to KDM2B, BCOR, BCORL1, and the accessory subunits SKP1 and USP7 (Gao et al., 2012). *BCOR* and *BCORL1* are recurrently mutated in hematologic malignancies, including acute and chronic leukemias. In acute myeloid leukemia (AML), *BCOR* and *BCORL1* mutations have been reported in 5-10% of patients and are associated with an antecedent diagnosis of myelodysplastic syndrome (MDS) and poor clinical outcomes after chemotherapy (Branford et al., 2018; Damm et al., 2013; Grossmann et al., 2011; Lindsley et al., 2015, 2017; Papaemmanuil et al., 2016; Terada et al., 2018). In chronic myeloid leukemia (CML), *BCOR* mutations have been linked to advanced disease and clinical resistance to BCR-ABL-targeted inhibitors (Branford et al., 2018). Leukemia-associated somatic *BCOR* and *BCORL1* mutations result in premature stop codons, but the mechanisms by which these mutations alter PRC1.1 complex assembly and function and contribute to leukemogenesis are unknown.

A major barrier to our understanding of the effect of *BCOR* and *BCORL1* mutations on PRC1.1 biology is a lack of information regarding *in vivo* subunit assembly, chromatin localization, and effects on target gene regulation. *In vitro* studies have suggested that BCOR/BCORL1 interaction with PCGF1 is required for subsequent assembly with KDM2B (Wong et al., 2016, 2020), but the consequences of BCOR/BCORL1 loss on complex integrity *in vivo* have not been shown. Studies have additionally ascribed distinct functions to other BCOR structural domains, such as the BCL6-binding and AF9-binding domains (Ghetu et al., 2008; Schmidt et al., 2020; Srinivasan et al., 2003), raising the possibility that somatic mutations may have heterogeneous effects on PRC1.1 activity, or could mediate effects on non-canonical PRC1.1 functions that are independent of H2A ubiquitination activity. Lastly, PRC1.1 has been shown to be broadly recruited to unmethylated CpG island promoters via the KDM2B CXXC domain (Farcas et al., 2012; He et al., 2013; Wu et al., 2013), but the subset of bound loci that are *bona fide* PRC1.1 functional targets has not been determined in leukemia.

Here, we combine genomic, biochemical and functional approaches to dissect the role of *BCOR* and *BCORL1* mutations in myeloid malignancies. We define the effect of individual subunit mutations on complex assembly, chromatin targeting, and target gene expression. These studies provide key insights into normal PRC1.1 function and the consequences of human disease-associated mutations, thereby uncovering a direct mechanistic link between epigenetic reprogramming and activation of signaling pathways that translates into a drug resistant phenotype.

## Results

### *BCOR* and *BCORL1* mutations cause protein loss or C-terminal protein truncations and significantly co-occur in myeloid malignancies

To define the frequency and spectrum of PRC1.1 mutations in AML, we performed targeted sequencing of genes encoding all nine PRC1.1 subunits in bone marrow samples from 433 AML patients. *BCOR* (26 of 433, 6%) and *BCORL1* (8 of 433, 1.8%) were most frequently mutated (Figure 1A). We identified one mutation in the complex-defining subunit *PCGF1* (0.2%), one mutation in *KDM2B* (0.2%), and no mutations in genes encoding other PRC1.1-specific accessory subunits (*SKP1*, *USP7*) or the pan-PRC1 enzymatic core (*RYBP*, *RING1*, *RNF2*) (Figure 1A, Table S1). These genetic findings indicate that selective alteration of PRC1.1 by *BCOR* and *BCORL1* mutations, but not alteration of PRC1 more globally, contributes to the pathogenesis of myeloid malignancies.

**Figure 1.**
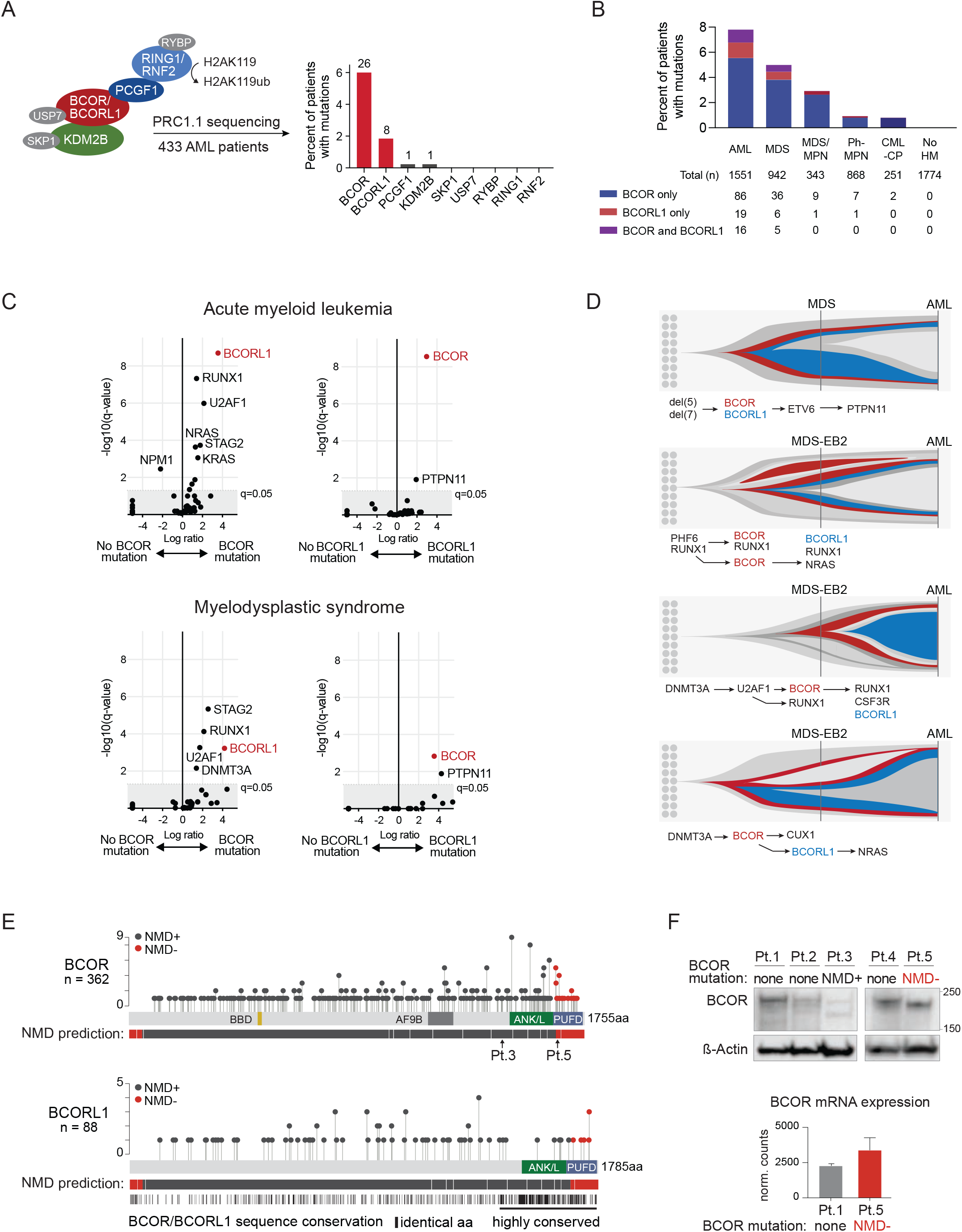
*BCOR* and *BCORL1* mutations cause protein loss or C-terminal protein truncations and significantly co-occur in myeloid malignancies. **(A)** Frequency of gene mutations in individual PRC1.1 subunits in a cohort of 433 AML patients. Number above bars indicate the number of patients with mutations. See also Table S1. **(B)** Bar plot representing disease distribution of *BCOR* and/or *BCORL1* mutations in a cohort of 3955 consecutive patients diagnosed with myeloid malignancies and 1774 individuals with hematologic abnormalities but no hematologic malignancy diagnosis. **(C)** Volcano plot representing associations of *BCOR* and *BCORL1* gene mutations in patients with AML (upper panel) or MDS (lower panel). The x axis shows the magnitude of association (log2 odds ratio), and the y axis the −log10 q-value. Each circle represents a mutated gene as indicated. Genes in the upper right quadrant significantly co-occurred. **(D)** Fish plots representing inferred clonal dynamics based on sequencing. See also Table S2. **(E)** Lollipop plot representing *BCOR* or *BCORL1* frameshift mutations. NMD sensitive (black circles; NMD+) and NMD insensitive (red circles; NMD-) mutations were defined using the prediction tool NMDetective-B (Lindeboom et al., 2019). PTCs with NMD scores <0.25 were considered NMD insensitive, PTCs with NMD scores >0.25 were considered NMD sensitive. Highly conserved regions of BCOR and BCORL1 were determined using Blastp (protein-protein Basic Local Alignment Search Tool) (Altschul et al., 1997). NMD: nonsense mediated mRNA decay. PTC: premature termination codon. **(F)** Western blot analysis (upper panel) of whole-cell lysates from BCOR^WT^ or *BCOR* mutated AML patient samples. The BCOR antibody RRID:AB_2716801 was used to detect BCOR protein levels (epitope: amino acids 1035-1230, (Gearhart et al., 2006)). Pt.3 harbors an NMD sensitive *BCOR* mutation (NMD+; p.S1439fs*44, predicted size: 161kDa). Pt.5 harbors an NMD insensitive *BCOR* mutation (NMD-; p.L1647fs*3, predicted size: 180kDa). *BCOR* gene expression levels (lower panel) represented as normalized DESeq2 counts in BCOR^WT^ (Pt.1) or BCOR^PUFD-Tr^ (Pt.5) primary AML patient samples (mean ± SD, n=2 technical duplicates for each condition).

In a separate cohort of 3955 consecutive patients diagnosed with myeloid malignancies at a single institution, we found that *BCOR* and *BCORL1* mutations were most common in AML (121/1551, 7.8%) and MDS (36/942, 5.0%), followed by MDS/MPN (10/343, 2.9%), Ph-MPN (8/868, 0.9%), and chronic phase CML (2/251, 0.8%) (Figure 1B). There were no *BCOR* or *BCORL1* mutations in 1774 consecutive patients with hematologic abnormalities but not found to have a hematologic malignancy. *BCOR* and *BCORL1* mutations significantly co-occurred in both MDS (LR=4.19, q<0.001) and AML (LR=3.57, q<0.0001), indicating that they may have cooperative effects in disease pathogenesis (Figure 1C). Consistent with this hypothesis, 45.7% (16/35) of AML patients with a *BCORL1* mutation also had a *BCOR* mutation, and serial samples from patients with concurrent *BCOR* and *BCORL1* mutations showed that the *BCORL1* mutation arose subclonally to a preexisting *BCOR* mutation (Figure 1D, Table S2).

Somatic *BCOR* (n=256) and *BCORL1* (n=90) mutations in this cohort were distributed across exons 4 to 15 and most (311 of 346, 89.9%) resulted in the introduction of premature termination codons (Table S3). Among all *BCOR* and *BCORL1* frameshift mutations, 90.8% (423/466) were predicted to cause reduced protein expression via nonsense mediated mRNA decay (NMD sensitive mutations, NMD+), and 9.2% (43/466) were predicted to cause stable expression of C-terminally truncated proteins that lacked part of the PCGF1 binding domain PUFD (NMD insensitive mutations, NMD-). The distribution of *BCOR* and *BCORL1* mutations in the two current cohorts, and in our previously published MDS (Lindsley et al., 2017) and AML (Lindsley et al., 2015) cohorts are shown in Figure 1E. Consistent with predictions, we detected expression of a PUFD truncated BCOR protein in primary AML blasts harboring an NMD insensitive mutation (Pt.5 BCOR p.L1647fs*3), whereas BCOR protein level was markedly reduced in a primary AML blasts harboring an NMD sensitive mutation (Pt.3 BCOR p.S1439fs*44) (Figure 1F). Disease-associated *BCOR* mutations thus result in either loss of protein expression or expression of C-terminal truncated protein that lacks an intact PUFD domain.

### *BCOR* and *BCORL1* mutations disrupt assembly of distinct PRC1.1 complexes by disrupting binding to PCGF1

To define the impact of *BCOR* and *BCORL1* mutations on PRC1.1 complex assembly and function, we generated isogenic K562 cell lines with frameshift *BCOR* and/or *BCORL1* mutations that resembled the spectrum of mutations that we observed in patients with myeloid malignancies (Figure 2A). Predicted NMD-sensitive *BCOR* mutations in exon 7, exon 9 or exon 10 resulted in loss of BCOR protein expression (BCOR^KO^) whereas predicted NMD-insensitive *BCOR* mutations in exon 14 caused expression of C-terminal truncated proteins (BCOR^PUFD-Tr^) (Figures 2B, S1A, and S1B). *BCORL1* mutations in exon 8 or exon 9 resulted in BCORL1 loss (BCORL1^KO^) (Figures 2C and S1C).

**Figure 2.**
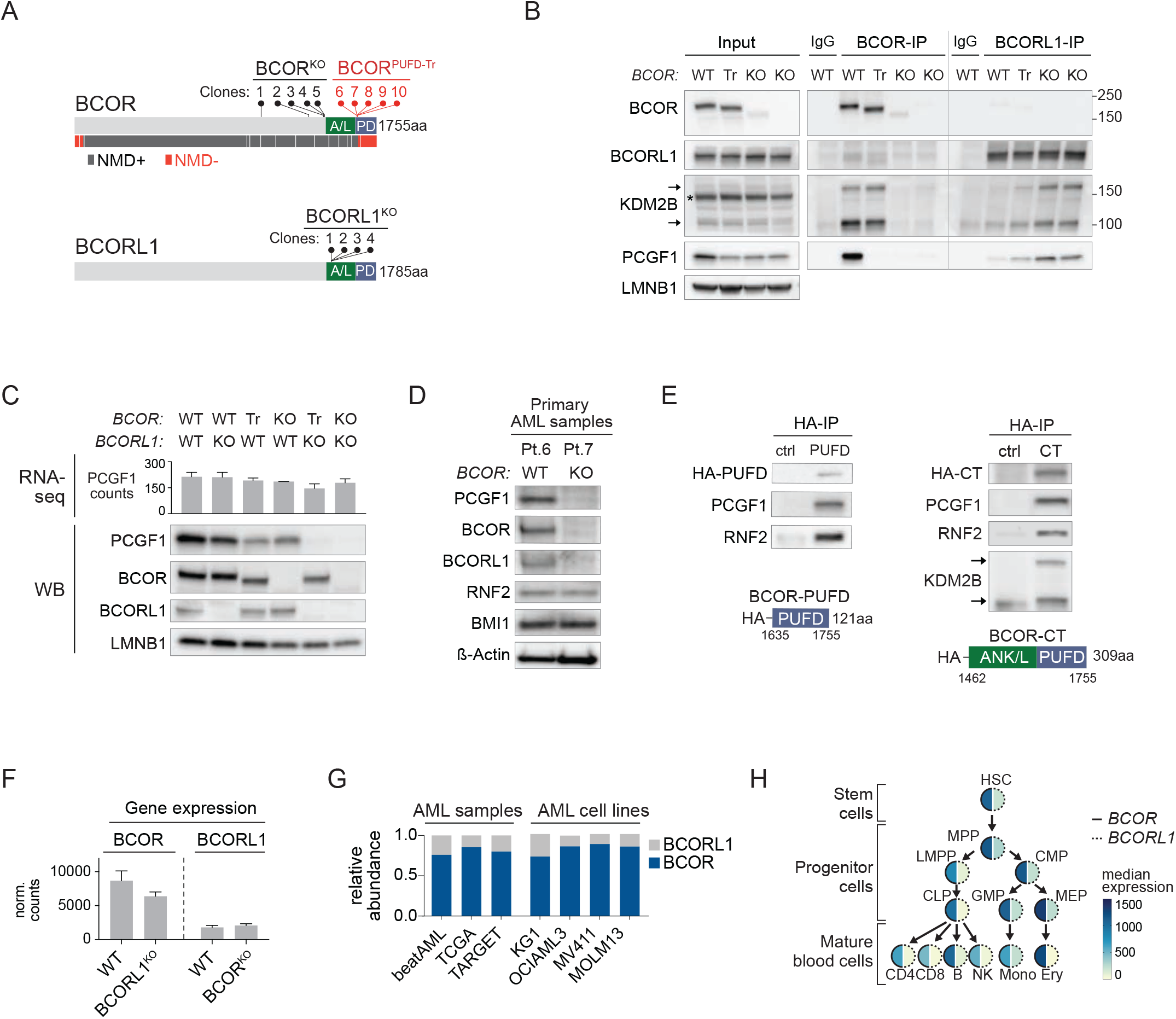
*BCOR* and *BCORL1* mutations disrupt assembly of distinct PRC1.1 complexes by disrupting binding to PCGF1. **(A)** Overview of BCOR and BCORL1 mutant isogenic cell lines (K562). Lollipops represent single cell clones with individual *BCOR* mutations (upper panel) or *BCORL1* mutations (lower panel). NMD sensitive (black circles; NMD+) and NMD insensitive (red circles; NMD-) mutations were defined using the prediction tool NMDetective-B (Lindeboom et al., 2019). PTCs with NMD scores <0.25 were considered NMD insensitive, PTCs with NMD scores >0.25 were considered NMD sensitive. A/L: ANK/L domains. PD: PUFD domains. See also Figures S1A and S1C. **(B)** BCOR- and BCORL1-Co-IP-western blot analysis in BCOR-WT (WT), BCOR^PUFD-Tr^ (Tr) or BCOR^KO^ (KO) K562 cells (nuclear extracts, endogenous proteins). KDM2B short (90kDa) and long (152kDa) isoforms are marked by arrows (asterisk indicates non-specific band). **(C)** *PCGF1* gene expression (DESeq2 normalized counts) in WT, BCORL1^KO^, BCOR^PUFD-Tr^, BCOR^KO^, BCOR^PUFD-Tr^/BCORL1^KO^ or BCOR^KO^/BCORL1^KO^ K562 cell lines represented as mean ± SD (upper panel). Lower panel shows corresponding western blot analysis of nuclear extracts. **(D)** Western blot analysis (whole cell lysates) of PRC1 subunits in BCOR^WT^ (Pt.6) or BCOR^KO^ (Pt.7, BCOR p.R1164*) AML patient cells. **(E)** Co-IP-western blot analysis of exogenously expressed BCOR HA-PUFD (1635-1755aa; left panel) or BCOR HA-CT (1462-1755aa; right panel) in BCOR^KO^/BCORL1^KO^ K562 cells. BCOR^KO^/BCORL1^KO^ cells transduced with a control vector were used as a negative control. **(F)** *BCOR* or *BCORL1* gene expression (DESeq2 normalized counts) in WT, BCORL1^KO^ or BCOR^KO^ K562 cells represented as mean ± SD. **(G)** Relative gene expression levels of *BCOR* and *BCORL1* in primary AML samples or AML cell lines as indicated. FPKM values of AML samples (beatAML n=507, TARGET n=532, LAML n=156) were obtained from the TCGA Research Network: https://www.cancer.gov/tcga. AML cell line RPKM values were downloaded from the Broad Institute Cancer Dependency Map (Broad Institute Cancer Dependency Map, 2018). **(H)** Median gene expression (DESeq2 normalized counts) of *BCOR* and *BCORL1* in hematopoietic stem/progenitor cells and mature lineages as indicated. RNA-seq counts of FACS-sorted cell lineages were obtained from Corces *et al*. (Corces et al., 2016).

Structural modeling has suggested that BCOR-PCGF1 and BCORL1-PCGF1 heterodimers are formed separately (Junco et al., 2013; Wong et al., 2016) but it is not known whether these heterodimers are recruited concurrently to individual PRC1.1 complexes or independently to distinct complexes *in vivo*. To distinguish between these possibilities, we performed reciprocal BCOR- and BCORL1-IP and western blot in WT cells (Figure 2B). We found that BCOR and BCORL1 were absent from IP of the reciprocal paralog, indicating that BCOR-PRC1.1 and BCORL1-PRC1.1 are mutually exclusive complexes. Further, BCORL1 interactions with PCGF1 and KDM2B were stable in the absence of BCOR, suggesting that BCOR-PRC1.1 and BCORL1-PRC1.1 complexes assemble independently.

Previously, it has been shown *in vitro* that BCORL1-PUFD mediates a direct interaction with PCGF1 that is required for subsequent interaction with KDM2B (Wong et al., 2016, 2020). In contrast, a recent study in hESCs showed that BCOR-PUFD was required to bind PCGF1 but not KDM2B (Wang et al., 2018). Using BCOR-Co-IP, we found that BCOR-PUFD truncations disrupted interactions with PCGF1 without affecting BCOR binding to KDM2B (Figure 2B). We validated our results by reciprocal PCGF1- and KDM2B-Co-IP (Figure S1D). Consistent with our findings in K562 cells, BCOR-PUFD truncations in MOLM14 cells disrupted binding to PCGF1 but not KDM2B (Figure S1E). These data indicate that BCOR-PUFD has an essential function in binding PCGF1 which is selectively disrupted even in patients with highly expressed C-terminal truncating *BCOR* mutations.

### BCOR and BCORL1 control PCGF1 protein levels via direct stabilizing interactions

PCGF1-RAWUL is reported to require direct interaction with BCOR/BCORL1-PUFD for stability *in vitro* (Junco et al., 2013), but the impact of this interaction on *in vivo* PRC1.1 complex assembly or on the stability of full length PCGF1 protein has not been shown. We therefore analyzed the effects of *BCOR* and *BCORL1* mutations on PCGF1 abundance in cells. In BCOR^KO^/BCORL1^KO^ conditions, we observed complete loss of PCGF1 protein, despite no significant differences in PCGF1 mRNA level (Figure 2C). In contrast, concurrent *BCOR/BCORL1* inactivation had no effect on the abundance of other PRC1.1-specific subunits (KDM2B, USP7, SKP1), pan-PRC1 enzymatic core subunits (RING1, RNF2, RYBP, CBX8), PCGF proteins (BMI1, PCGF3, PCGF5), or PRC2 (EZH2) (Figure S1F). PCGF1 protein level was decreased in primary AML samples with *BCOR* mutations (Figures 2D and S1G).

We observed a similar selective effect on PCGF1 protein level in BCOR^PUFD-Tr^/BCORL1^KO^ cells (Figure 2C), indicating that BCOR protein lacking the PUFD is unable to stabilize PCGF1. To test whether the C-terminal PCGF1-binding domains of BCOR were alone sufficient to restore BCOR-PCGF1 interactions and PCGF1 protein stability *in vivo*, we expressed HA-tagged BCOR-PUFD or BCOR-CT domains in BCOR^KO^/BCORL1^KO^ cells (Figure 2E). Expression of BCOR-PUFD was sufficient to form a stable heterodimer with PCGF1 and restore interactions with RNF2 (Figure 2E). In line with previous findings (Wang et al., 2018; Wong et al., 2020), the BCOR-CT domain, which contains the ANK/linker and the PUFD domains, similarly restored the stable interaction with PCGF1, but also bound to KDM2B (Figures 2E and S1H). Thus, the PUFD domain was necessary and sufficient to bind and stabilize PCGF1 protein, while the ANK/linker was required for co-recruitment of KDM2B.

Sole disruption of BCOR (BCOR^KO^ or BCOR^PUFD-Tr^) caused only partial reduction of PCGF1 protein levels and sole disruption of BCORL1 (BCORL1^KO^) had no impact on PCGF1 protein abundance (Figure 2C), indicating that persistent expression of paralogous BCORL1 or BCOR compensates for BCOR or BCORL1 loss, respectively. We hypothesized that the differential impact of BCOR versus BCORL1 loss on total PCGF1 protein level was due to differences in the abundance of paralog expression at baseline. Consistent with this hypothesis, BCOR was expressed more highly than BCORL1 in WT K562 cells (Figure 2F). More broadly, BCOR was the predominantly expressed paralog in other BCOR/BCORL1^WT^ AML cell lines (KG1, OCIAML3, MV4-11 and MOLM13), in primary AML samples, and throughout normal hematopoiesis, including stem and progenitor cells, and mature blood cells across all lineages (Figures 2G and 2H). Concordant with the higher BCOR expression levels, BCOR contributed to the majority of PCGF1 interactions in Co-IP analysis (Figure S1I). We further observed no compensatory increase of BCORL1 gene expression or protein levels in BCOR mutant cells (Figure 2F). Together, our data indicate that PCGF1 abundance is directly linked to the composite level of BCOR and BCORL1 expression levels and that reduction of PCGF1 levels in single BCOR or BCORL1 mutant cells is proportional to the baseline expression levels of the remaining paralog.

### Loss of BCOR-PCGF1 or BCORL1-PCGF1 interaction disrupts complex assembly by uncoupling the enzymatic core from KDM2B

We showed that a shared effect of BCOR and BCORL1 inactivation is to destabilize PCGF1 protein levels. Therefore, we next sought to define the impact of direct PCGF1 loss on PRC1.1 complex assembly. To generate isogenic PCGF1^KO^ cell lines, we introduced mutations in the PCGF1-RING domain or in a region that was previously identified as an essential contact site for PCGF1-PUFD interactions (Figure 3A) (Wong et al., 2016). PCGF1 loss did not impair protein stability of BCOR or BCORL1 (Figure 3A).

**Figure 3.**
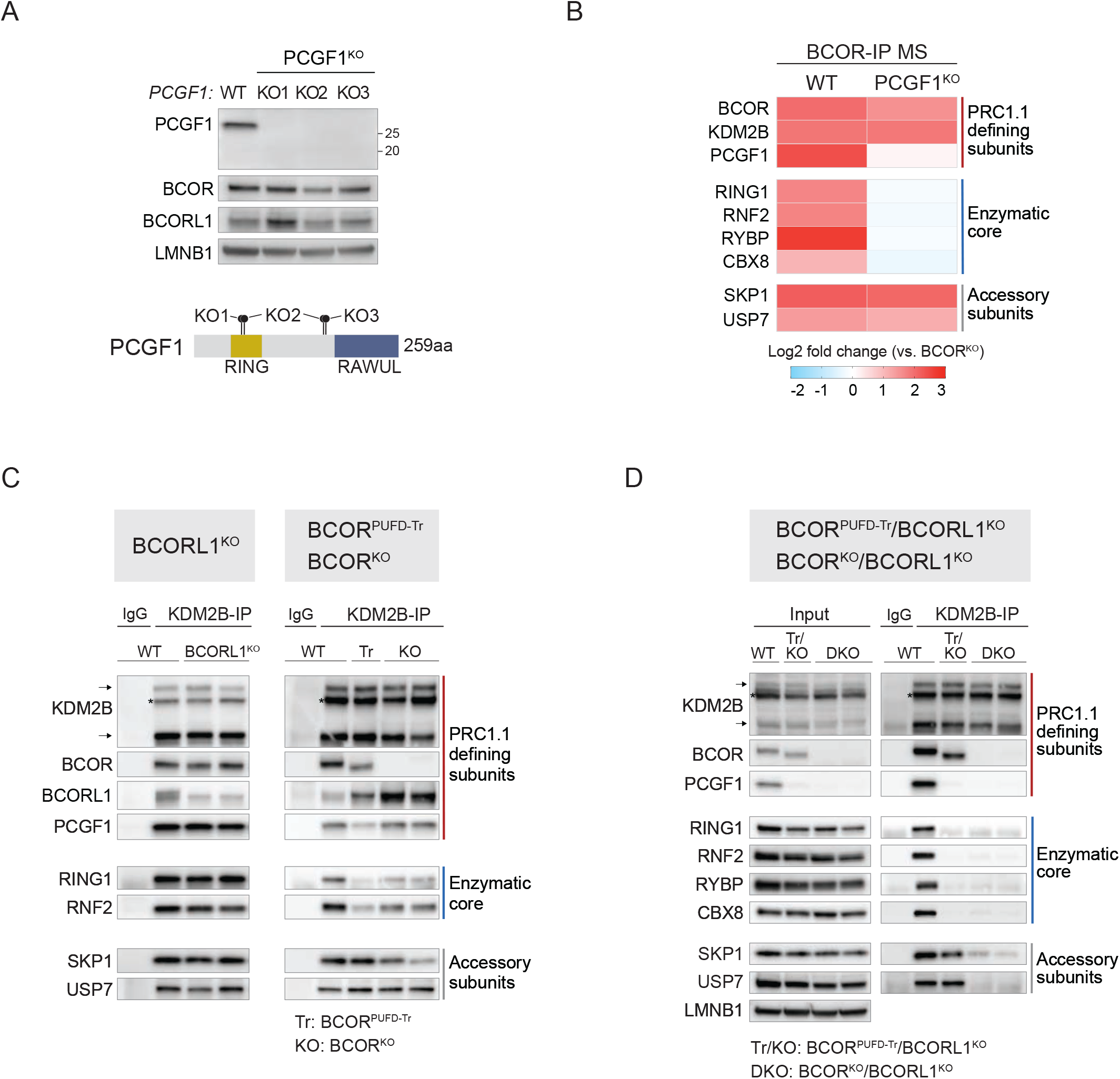
Loss of BCOR-PCGF1 or BCORL1-PCGF1 interaction disrupts complex assembly by uncoupling the enzymatic core from KDM2B. **(A)** Western blot analysis of nuclear extracts from WT or PCGF1^KO^ K562 cells. PCGF1 mutations are illustrated in the schematic below. KO1: PCGF1-p.I60fs*. KO2: PCGF1-p.(I60fs*;p.Y163fs*). KO3: PCGF1-p.Y163del. The PCGF1 antibody RRID:AB_2721914 was used to detect PCGF1 protein levels (epitope: amino acids 128-247). **(B)** Enrichment of PRC1.1 subunits in BCOR-IP mass spectrometry analysis in WT or PCGF1^KO^ K562 cells represented as log2 fold change compared to BCOR^KO^ control. n = 2 (process replicates for each condition). See also Figures S2A-C. **(C)** KDM2B-Co-IP-western blot analysis (elution fractions, nuclear extracts) in WT or BCORL1^KO^ K562 cells (left panel), and in WT, BCOR^PUFD-Tr^ (Tr) or BCOR^KO^ (KO) K562 cells (right panel). KDM2B short (90kDa) and long (152kDa) isoforms are marked by arrows (asterisk indicates non-specific band). **(D)** KDM2B-Co-IP-western blot analysis (nuclear extracts) in WT, BCOR^PUFD-Tr^/BCORL1^KO^ (Tr/KO) or BCOR^KO^/BCORL1^KO^ (DKO) K562 cells. KDM2B short (90kDa) and long (152kDa) isoforms are marked by arrows (asterisk indicates non-specific band).

To identify BCOR-interacting proteins in the presence or absence of PCGF1, we immunoprecipitated BCOR and analyzed the complex by mass spectrometry (MS) in WT or PCGF1^KO^ cells. In WT cells, the main interaction partners of BCOR were PRC1.1-defining and accessory subunits (PCGF1, KDM2B, USP7, and SKP1) and pan-PRC1 enzymatic core subunits (RING1/RNF2, RYBP, and CBX8) (Figures 3B, S2A, and S2B). We validated these results in K562 cells using IP-western blot (Figure S2C) and confirmed BCOR interactions with core PRC1.1 subunits in primary AML patient samples and in OCI-AML3 and KG1 AML cell lines (Figure S2D). BCORL1 interactions with PRC1.1-specific and pan-PRC1 subunits mirrored those of BCOR (Figure S2E). In PCGF1^KO^ cells, we observed loss of BCOR binding to the enzymatic core, including RING1, RNF2, RYBP, and CBX8 (Figures 3B and S2A-S2C).

However, the absence of PCGF1 did not impair BCOR interactions with KDM2B, SKP1, and USP7, indicating that PCGF1 is essential for coupling the enzymatic core to the complex but not for mediating BCOR-KDM2B interactions.

To determine whether indirect loss of PCGF1 protein via *BCOR* and *BCORL1* mutations had the same effect on PRC1.1 assembly as direct loss of PCGF1 via gene deletion, we performed KDM2B-Co-IP in BCORL1^KO^, BCOR^PUFD-Tr^, BCOR^KO^, BCOR^PUFD-Tr^/BCORL1^KO^, and BCOR^KO^/BCORL1^KO^ cells. Alone, neither *BCOR* nor *BCORL1* mutations caused complete disruption of KDM2B interactions with PRC1.1 subunits (Figure 3C), consistent with our observation that BCOR-PRC1.1 and BCORL1-PRC1.1 form independent complexes with a redundant adaptor function in PRC1.1 assembly. We observed similar results in BCOR^PUFD-Tr^ MOLM14 cells (Figure S2F). Concurrent disruption of BCOR and BCORL1 resulted in complete uncoupling of the PRC1 enzymatic core from KDM2B, reflected by loss of RING1, RNF2, RYBP, and CBX8 binding (Figure 3D). This finding was identical in cells lacking BCOR protein expression (BCOR^KO^/BCORL1^KO^) and cells expressing the PUFD-truncated BCOR (BCOR^PUFD-Tr^/BCORL1^KO^), further confirming the specific and absolute requirement for BCOR-PUFD in PRC1.1 complex assembly. To validate this finding, we complemented BCOR^KO^/BCORL1^KO^ cells via exogenous expression of full-length BCOR and showed restoration of PCGF1 protein levels and KDM2B interactions with the enzymatic core subunits (Figure S2G).

### KDM2B is essential to recruit BCOR-PRC1.1 and BCORL1-PRC1.1 to unmethylated CpG island promoters

To determine the localization of BCOR- and BCORL1-PRC1.1 complexes at chromatin, we performed BCOR, BCORL1, KDM2B, and RNF2 ChIP-seq in K562 cells. BCOR and BCORL1 bound predominantly to promoter regions (Figures 4A and S3A) and the distributions of BCOR and BCORL1 binding at transcription start sites (TSS) were significantly correlated (R^2^=0.7375, p<0.0001; Figure 4B). BCORL1 binding was unchanged in BCOR^KO^ cells compared with WT cells (Figures 4A and 4B) and BCOR- and BCORL1-bound genes were co-occupied by the PRC1.1 subunits KDM2B and RNF2 (Figure 4A). Together, these results indicate that BCOR-PRC1.1 and BCORL1-PRC1.1 complexes are independently recruited to a shared set of target loci.

**Figure 4.**
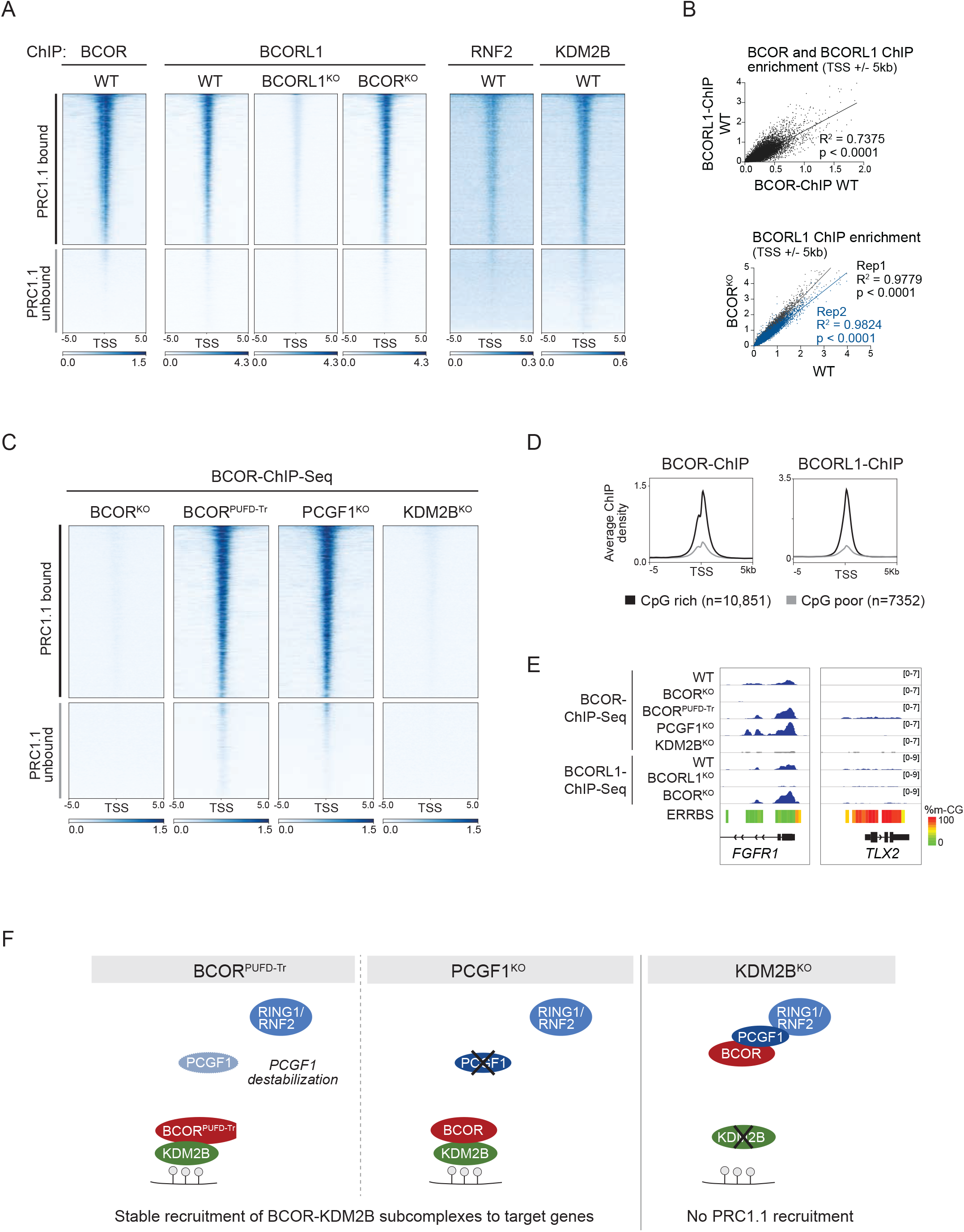
KDM2B is essential to recruit BCOR-PRC1.1 and BCORL1-PRC1.1 to unmethylated CpG island promoters. **(A)** Heatmaps represent BCOR-, BCORL1-, KDM2B-, and RNF2-ChIP enrichment at transcription start sites (TSS) +/- 5kb in WT, BCORL1^KO^, or BCOR^KO^ K562 cells. BCOR-ChIP signal scores in WT K562 cells were used to predefine PRC1.1 bound (n=11,934) and PRC1.1 unbound (n=6,639) promoters. All heatmaps are sorted and ranked based on BCOR-ChIP enrichment in WT cells. **(B)** Scatterplots showing the relationship between BCORL1- and BCOR-ChIP-seq signals at TSS +/- 5kb in WT K562 cells (upper panel). Scatterplots showing the relationship between BCORL1-ChIP-seq signals in BCOR^KO^ and WT K562 cells (lower panel). Biological replicates of BCOR^KO^ conditions are plotted in black (Rep1) or in blue (Rep2). Pearson correlation coefficients and p values (two-tailed) as indicated. **(C)** Heatmaps of BCOR-ChIP-seq enrichment at PRC1.1 bound (n=11,934) and PRC1.1 unbound (n=6,639) gene promoters in BCOR^KO^, BCOR^PUFD-Tr^, PCGF1^KO^, or KDM2B^KO^ cells. All heatmaps are sorted and ranked based on BCOR-ChIP signals in WT cells (Figure 4A). **(D)** Average density profiles of BCOR-ChIP and BCORL1-ChIP signals in WT K562 cells at CpG rich promoters (n=10,851) or CpG poor promoters (n=7352). See also Figure S3F. **(E)** Genomic coverage data tracks for BCOR-ChIP-seq and BCORL1-ChIP-seq at unmethylated (*FGFR1*) or methylated (*TLX2*) CpG island promoters. Methylation percentages of CpG sites are indicated by the color gradient. **(F)** Schematic of PRC1.1 recruitment in BCOR^PUFD-Tr^, PCGF1^KO^, and KDM2B^KO^ conditions. BCOR is stably recruited to target genes in BCOR^PUFD-Tr^ and PCGF1^KO^ conditions but not in KDM2B^KO^ cells, indicating that KDM2B but not the enzymatic core is required for PRC1.1 recruitment in leukemia cells.

In human ESCs, PRC1.1 binding to most target genes has been reported to be dependent on RING1/RNF2 (Wang et al., 2018), while in mouse ESCs PRC1.1 recruitment is largely mediated by KDM2B (Farcas et al., 2012; He et al., 2013; Wu et al., 2013). This suggests that the recruitment mechanism of PRC1.1 may be cell type dependent. To determine the differential requirement for the PRC1 enzymatic core and KDM2B in PRC1.1 complex recruitment in human leukemia cells, we performed BCOR-ChIP-seq in WT cells or cells deficient for different PRC1.1 subunits. In both BCOR^PUFD-Tr^ and PCGF1^KO^ cells, where BCOR is selectively uncoupled from the enzymatic core while retaining binding to KDM2B, the number of BCOR peaks was similar compared to WT (Figures 4C, S3A and S3C) and the distribution of BCOR binding at TSS was significantly correlated with BCOR enrichment in WT (WT vs. BCOR^PUFD-Tr^, R^2^ = 0.8949, p<0.0001 and WT vs. PCGF1^KO^, R^2^ = 0.9413, p<0.0001; Figure S3B). We validated this finding in primary patient samples, confirming that truncated BCOR maintained binding to target genes (Figure S3D).

To test the requirement for KDM2B in PRC1.1 targeting, we generated isogenic KDM2B^KO^ cell lines by introducing frameshift mutations upstream of the DNA-binding CXXC domain that resulted in loss of both long and short isoforms. In these KDM2B^KO^ conditions, BCOR retained normal interactions with PRC1.1-specific subunits (USP7 and PCGF1) and the PRC1 enzymatic core, but lost KDM2B-dependent binding to SKP1 (Figure S3E). In the absence of KDM2B, we observed global loss of BCOR chromatin binding that was equivalent to negative control BCOR^KO^ conditions (Figures 4C, 4E, and S3C), indicating that PRC1.1 targeting to chromatin is absolutely dependent on KDM2B. KDM2B has been reported to bind to DNA via its CXXC zinc-finger domain, which recognizes unmethylated CpG islands (CGIs) (Farcas et al., 2012; He et al., 2013; Wu et al., 2013). Consistent with a KDM2B-dependent recruitment mechanism, we found that BCOR and BCORL1 binding was enriched at CGI promoters and positively correlated with CpG abundance at CGI promoters in K562 cells (Figures 4D and S3F), as well as in in primary AML patient samples and in KG1, OCI-AML3, and MV411 leukemia cell lines (Figures S3G and S3H). Using enhanced reduced representation bisulfite sequencing (ERRBS) in WT K562 cells, we found that BCOR and BCORL1 occupancy was negatively correlated with DNA methylation levels at CGIs (Figures 4E and S3I). Together, these results indicate that KDM2B, but not PCGF1 or the enzymatic core, is essential to recruit BCOR- and BCORL1-PRC1.1 to unmethylated CpG islands in leukemia (Figure 4F).

### Recruitment of the PRC1.1 enzymatic core is required for target gene repression

We next surveyed the chromatin state of PRC1.1-bound and PRC1.1-unbound genes using H3K27ac, H3K4me3, H3K27me3, and H2AK119ub ChIP-seq and ATAC-seq. Among PRC1.1-bound genes, 78.2% (9336 of 11,934) had an active chromatin state reflected by high levels of H3K27ac and H3K4me3, high chromatin accessibility, and high baseline gene expression (Figures 5A and S4A). 21.8% (2598 of 11,934) of PRC1.1-bound genes had a repressed chromatin state with high H3K27me3, high H2AK119ub, and low or no gene expression (Figures 5A and S4A). In contrast, PRC1.1-unbound genes (n=6269) largely had an inactive chromatin state, associated with compacted chromatin and no gene expression (Figures 5A and S4A). We observed a similar distribution of BCOR binding in KG1, OCI-AML3, and MV4-11 cell lines and in primary AML patient cells (Figures S4A-S4D).

**Figure 5.**
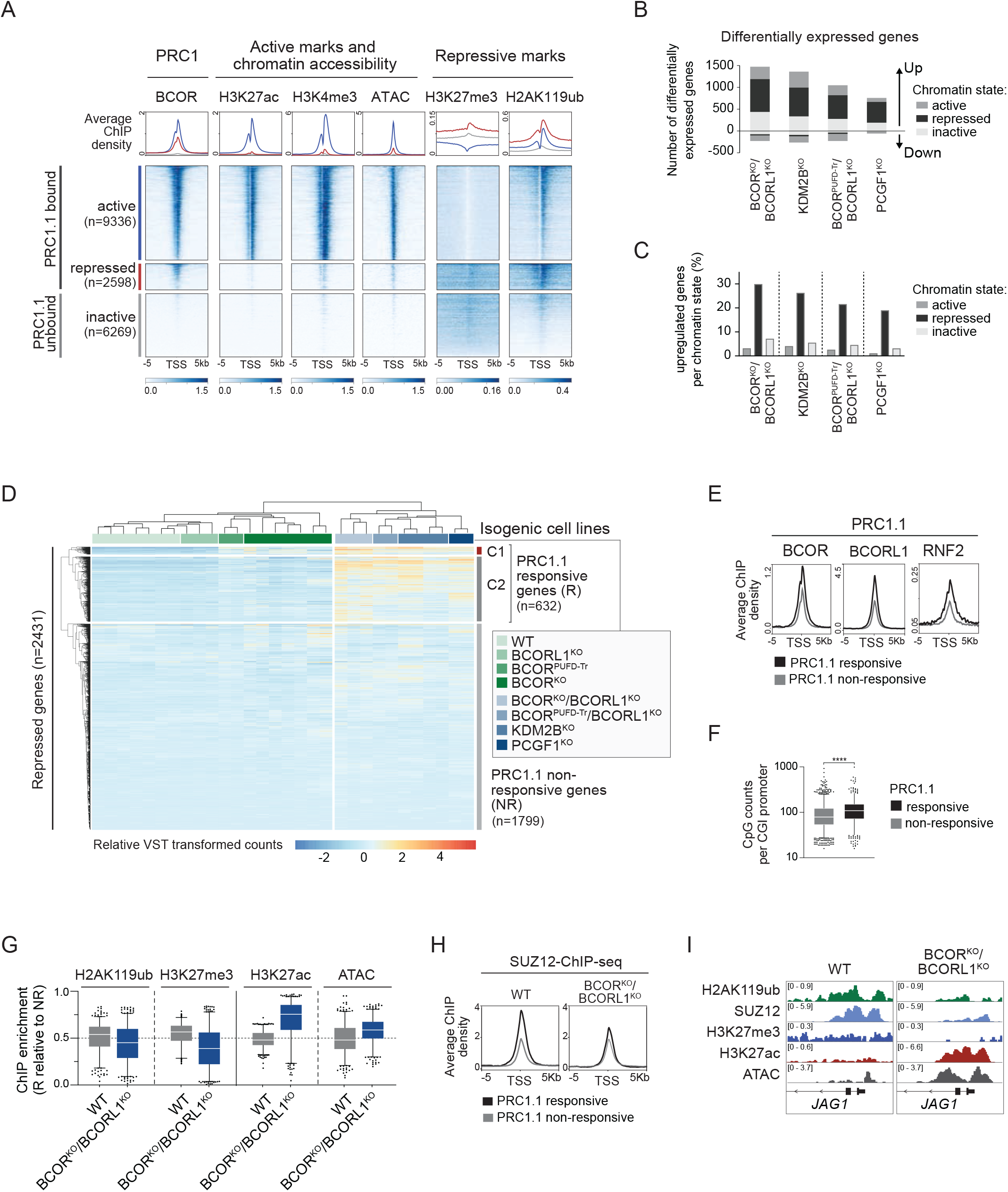
Recruitment of the PRC1.1 enzymatic core is required for target gene repression. **(A)** Heatmaps and profile plots of ChIP-seq enrichment of BCOR, histone modifications, and chromatin accessibility signals (ATAC) as indicated. BCOR-ChIP signal scores in K562 WT cells were used to predefine PRC1.1 bound promoters (n=11,934). BCOR- and H3K27ac-ChIP signal scores in K562 WT cells were used to define active, repressed, and inactive gene promoters. K562 H3K4me3-ChIP-seq data was downloaded from the Sequence Read Archive (SRA; SRR341173). The remaining datasets were generated in this study. See also Figure S4A. **(B)** Number of differentially expressed genes in PRC1.1 mutant compared to WT K562 cells (fold change >2, padj <0.05). Differentially expressed genes were subdivided based on their chromatin states defined in Figure 5A. **(C)** Bar diagram represents the percentages of upregulated genes in PRC1.1 mutant cells relative to the total number of genes in each chromatin state. **(D)** Unsupervised hierarchical clustering analysis of expression levels of PRC1.1 bound repressed genes in WT (n=7) or PRC1.1 mutant K562 cells (BCORL1^KO^ (n=3), BCOR^PUFD-Tr^ (n=2), BCOR^KO^ (n=7), BCOR^KO^/BCORL1^KO^ (n=3), BCOR^PUFD-Tr^/BCORL1^KO^ (n=2), KDM2B^KO^ (n=4), PCGF1^KO^ (n=2)). Genes clustered into PRC1.1 responsive (n=632) or PRC1.1 non-responsive genes (n=1799). PRC1.1 responsive genes clustered into C1 (n=65) or C2 (n=567) genes. Gene expression levels are shown as relative VST transformed values. See also Figure S4G. **(E)** Average density profiles of BCOR-, BCORL1- and RNF2-ChIP signals in WT K562 cells at PRC1.1 responsive (n=632) or PRC1.1 non-responsive genes (n=1799). **(F)** Number of CpG counts at PRC1.1 responsive (n=632) or non-responsive CGI promoters (n=1799). ****p<0.0001, unpaired t test. The line in the middle of the box represents the median, box edges represent the 25th and 75th percentiles, and whiskers show 5th and 95th percentiles. **(G)** Box and whisker plots representing histone ChIP- and ATAC-seq signals at PRC1.1 responsive genes (n=632) normalized to the signals at PRC1.1 non-responsive genes (n=1799) in WT or BCOR^KO^/BCORL1^KO^ cells. The line in the middle of the box represents the median, box edges represent the 25th and 75th percentiles, and whiskers show 2.5th and 97.5th percentiles. **(H)** Average density profiles of SUZ12-ChIP signals in WT or BCOR^KO^/BCORL1^KO^ K562 cells at PRC1.1 responsive (n=632) or PRC1.1 non-responsive genes (n=1799). **(I)** ChIP-seq and ATAC-seq coverage data tracks of the PRC1.1 responsive gene *JAG1* in WT or BCOR^KO^/BCORL1^KO^ K562 cells.

PRC1 chromatin binding alone does not directly correlate with regulatory activity (Farcas et al., 2012; Klose et al., 2013). Therefore, we determined the set of PRC1.1 functional targets by analyzing the impact of complete PRC1.1 inactivation on global gene expression compared with PRC1.1 WT cells. We found that the predominant consequence was gene upregulation, irrespective of the mechanism of complex inactivation, including complete PRC1.1 loss (BCOR^KO^/BCORL1^KO^: 85.6%, 1481/1731), disruption of PRC1.1 recruitment to chromatin (KDM2B^KO^: 83.3%, 1380/1656), and selective uncoupling of the PRC1 enzymatic core (BCOR^PUFD-Tr^/BCORL1^KO^: 81.8%, 1060/1296 and PCGF1^KO^: 92.3%, 756/819) (Figure 5B). The proportion of differentially expressed genes was highest among genes with repressed chromatin state at baseline, and genes with active and inactive chromatin state exhibited relatively small changes (Figures 5B and S4E). Indeed, among the set of repressed PRC1.1-bound genes, we observed a significant gene upregulation in BCOR^KO^/BCORL1^KO^ (29.8%), KDM2B^KO^ (26.2%), BCOR^PUFD-Tr^/BCORL1^KO^ (21.5%), and PCGF1^KO^ (18.9%) (Figure 5C), and there was high overlap of upregulated repressed genes across all PRC1.1 mutant conditions (Figure S4F). Together, these results indicate that PRC1.1 disruption causes selective derepression of target genes.

To identify a set of PRC1.1 target genes among genes with repressed chromatin at baseline, we performed unsupervised hierarchical clustering of gene expression in WT and all PRC1.1 mutant cells. We identified two main gene clusters: PRC1.1 responsive genes (n=632) and PRC1.1 non-responsive genes (n=1799) (Figure 5D). Gene expression levels of PRC1.1 responsive genes were highly increased in all PRC1.1 inactive conditions (BCOR^PUFD-Tr^/BCORL1^KO^, BCOR^KO^/BCORL1^KO^, PCGF1^KO^, KDM2B^KO^) compared with WT control, whereas PRC1.1 non-responsive genes showed similar expression levels in WT and PRC1.1 mutant cells (Figure S4G). To identify parameters that were associated with PRC1.1 responsiveness we assessed PRC1.1 occupancy and histone modifications at PRC1.1 responsive or non-responsive genes. PRC1.1 responsive genes showed higher levels of H2AK119ub and H3K27me3 than PRC1.1 non-responsive genes (Figure S4H). We further observed stronger binding of BCOR, BCORL1, and RNF2 at responsive compared to non-responsive genes which correlated with higher CpG levels at CGI promoters (Figures 5E and 5F), indicating that PRC1.1 was required to maintain repression of a defined set of CpG rich target genes.

Recent studies highlighted that H2AK119ub deposition by non-canonical PRC1 was essential to recruit PRC2 and to maintain repression of target genes (Blackledge et al., 2020; Fursova et al., 2019; Tamburri et al., 2020). We hypothesized that PRC1.1 loss disrupted H2AK119ub deposition resulting in epigenetic activation of PRC1.1 responsive genes. To test this hypothesis, we performed H2AK119ub, H3K27me3, H3K27ac, and ATAC-seq in BCOR^KO^/BCORL1^KO^ cells. We observed decreased H2AK119ub and H3K27me3 levels at PRC1.1 responsive genes in BCOR^KO^/BCORL1^KO^ cells compared to WT cells while H3K27ac levels and chromatin accessibility were increased (Figures 5G and 5I). PRC1.1 disruption and decreased H2AK119ub deposition at PRC1.1 responsive genes were further associated with reduced PRC2 occupancy (Figures 5H and 5I) whereas global levels of H2AK119ub or PRC2 were not impaired (Figure S4I). These data suggest that PRC1.1 enzymatic activity maintains active repression of specific target genes by H2AK119ub deposition and stable recruitment of PRC2.

### PRC1.1 regulates cell signaling programs in human leukemia

To identify PRC1.1 dependent transcription programs in leukemia cells, we performed gene set enrichment analysis of differentially expressed genes in BCOR^KO^/BCORL1^KO^ cells compared with BCOR^WT^/BCORL1^WT^ cells. Differentially expressed genes in BCOR^KO^/BCORL1^KO^ conditions were enriched for signaling pathways regulating stem cell pluripotency (KEGG hsa04550) including FGF/FGFR signaling, PI3K-Akt signaling and TGF-ß signaling pathways (Figures 6A, 6B, S5A, and S5B), consistent with a role of PRC1.1 in regulating stem cell signaling programs. We further observed enrichment of signaling pathways involved in cancer (KEGG hsa05200) such as RAS and MAPK signaling pathways (Figures 6A, S5A, and S5B).

**Figure 6.**
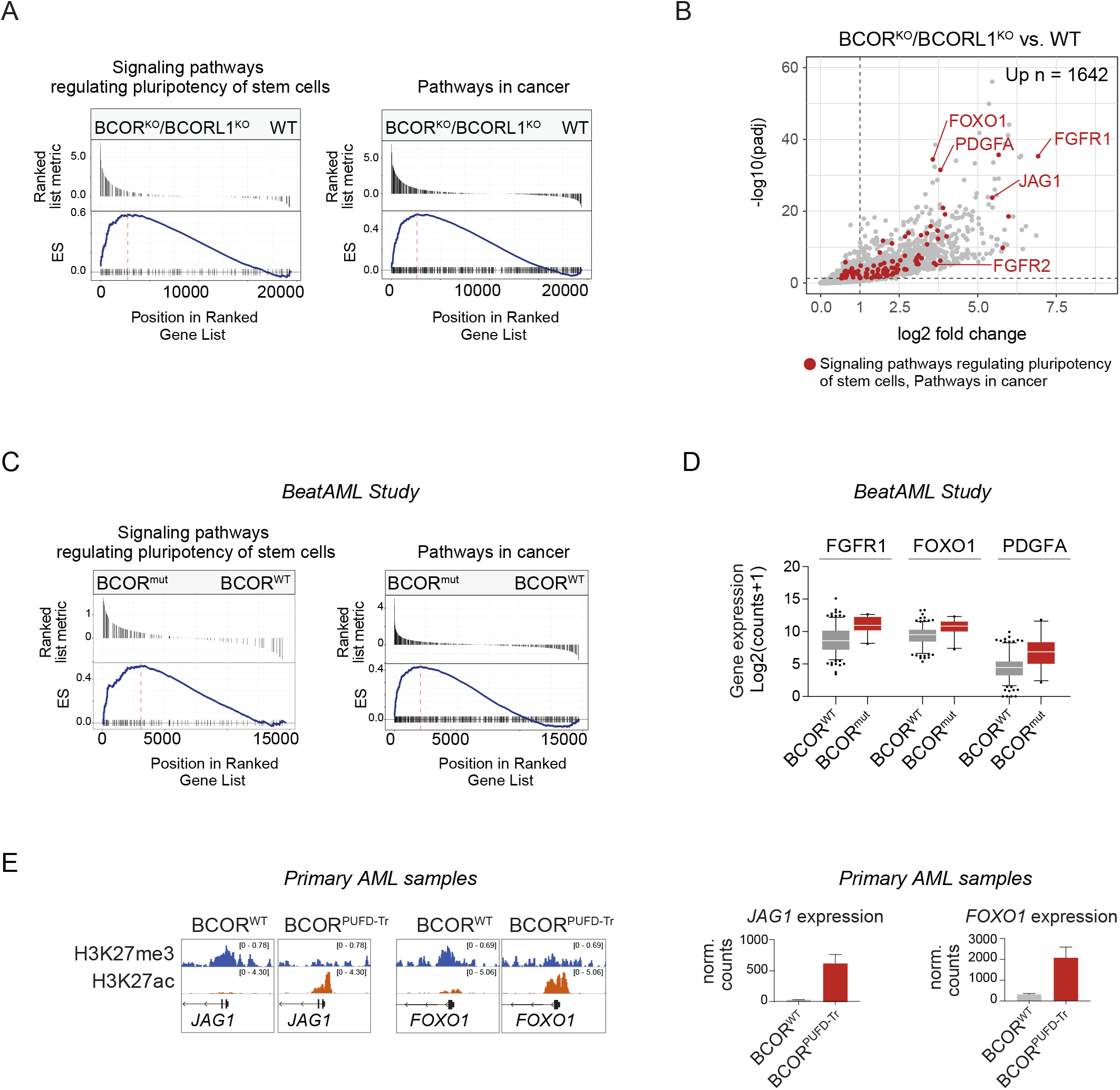
PRC1.1 regulates cell signaling programs in human leukemia. **(A)** KEGG gene set enrichment analysis (GSEA) of differentially expressed genes in BCOR^KO^/BCORL1^KO^ compared to WT K562 cells. The top plot represents the magnitude of log2 fold changes for each gene. The bottom plot indicates the enrichment score (ES). See also Figure S5A. **(B)** Volcano plot representing gene expression changes of upregulated genes in BCOR^KO^/BCORL1^KO^ compared to WT K562 cells. Genes represented in KEGG Signaling pathways regulating pluripotency of stem cells (hsa04550) or KEGG Pathways in cancer (hsa05200) are highlighted in red. Log2 fold changes of 1 and padj values of 0.05 are depicted by dotted lines. 1642 genes were significantly upregulated more than 2-fold in BCOR^KO^/BCORL1^KO^ cells compared to WT control. **(C)** KEGG gene set enrichment analysis of differentially expressed genes in BCOR^mut^ (n=22) compared to BCOR^WT^ (n=216) AML patient samples from the beatAML study (Tyner et al., 2018). The top plot represents the magnitude of log2 fold changes for each gene. The bottom plot indicates the enrichment score (ES). See also Figure S5C. **(D)** Box and whisker plots representing expression levels (log2 transformed DESeq2 normalized counts) in BCOR^WT^ (n=216) or BCOR^mut^ (n=22) primary AML samples (Tyner et al., 2018). The line in the middle of the box represents the median, box edges represent the 25th and 75th percentiles, and whiskers show 5th and 95th percentiles. **(E)** ChIP-seq coverage data tracks of PRC1.1 target genes *JAG1* and *FOXO1* in BCOR^WT^ (Pt.1) or BCOR^PUFD-Tr^ (Pt.5) cells (left panel). Corresponding gene expression levels of *JAG1* and *FOXO1* (DESeq2 normalized counts, n=2, technical replicates; right panel).

To validate our results in primary patient samples, we performed differential gene expression analysis in samples with (n=22) and without (n=216) *BCOR* mutations from the Beat AML study (Tyner et al., 2018). Differentially expressed genes in BCOR mutated samples were enriched for regulators of stem cell pluripotency and cancer signaling pathways with higher gene expression levels of FGFR1, PDGFA, and FOXO1 (Figures 6C, 6D, S5C, and S5D), similar to what we observed in BCOR mutated cell line models.

Using H3K27me3 and H2AK119ub ChIP-seq and RNA-seq analysis, we found that the PRC1.1 target genes JAG1 and FOXO1 had a repressed chromatin state and were lowly expressed in primary BCOR^WT^ AML blasts, whereas these genes were associated with an active chromatin state in primary BCOR^PUFD-Tr^ AML blasts (Figures 6E and S5E). Altogether, these data indicate that PRC1.1 inactivation in BCOR mutated AML results in aberrant expression of stem cell signaling programs via epigenetic derepression of target genes.

### BCOR^KO^ causes partial PRC1.1 reduction and activates a set of highly responsive target genes

In patients with myeloid malignancies, *BCOR* mutations are most commonly found without concurrent *BCORL1* mutations, indicating that partial PRC1.1 disruption is sufficient to drive clonal advantage. We hypothesized that a subset of functionally relevant PRC1.1 target genes are particularly sensitive to quantitative decreases in PRC1.1 abundance. We therefore compared the effect of single BCOR^PUFD-Tr^ and BCOR^KO^ conditions on gene expression with that of double mutant conditions (BCOR^PUFD-Tr^/BCORL1^KO^ and BCOR^KO^/BCORL1^KO^) and found that a subset of genes that were upregulated in double BCOR/BCORL1 mutant conditions were also induced in single BCOR mutants. (Figure 7A and S6A).

**Figure 7.**
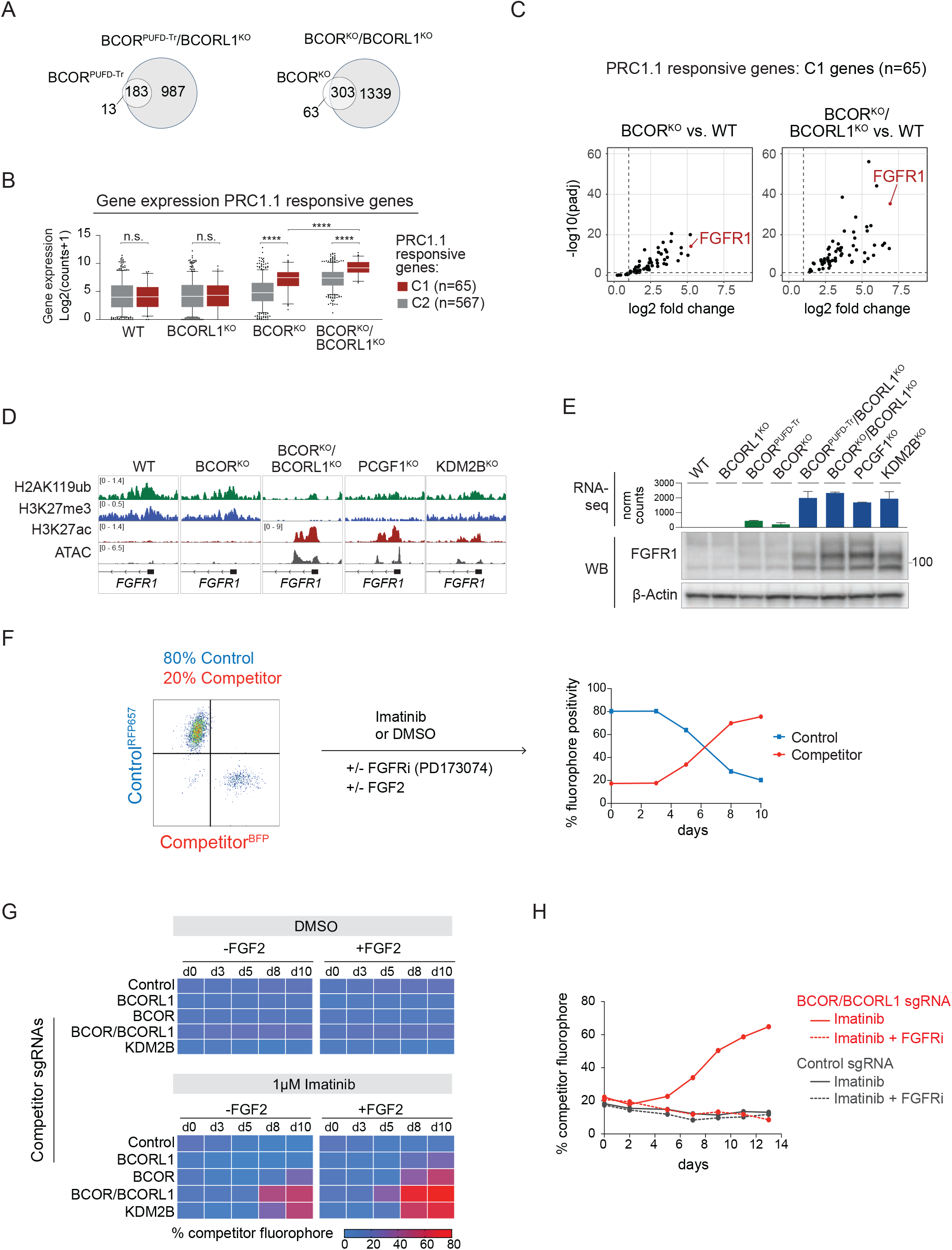
PRC1.1 mutations drive aberrant FGFR1 activation which mediates resistance to targeted therapy. **(A)** Venn diagrams represent the overlaps of significantly upregulated genes in PRC1.1 mutant cells vs. WT control. **(B)** Box and whisker plots representing expression levels (log2 transformed DESeq2 normalized counts) of C1 (n=65) and C2 (n=567) genes in WT, BCORL1^KO^, BCOR^KO^ or BCOR/BCORL1^KO^ cells. C1 and C2 genes were defined using unsupervised hierarchical clustering analysis of expression levels of PRC1.1 responsive genes in WT or PRC1.1 mutant cells shown in Figure 5C. The line in the middle of the box represents the median, box edges represent the 25th and 75th percentiles, and whiskers show 5th and 95th percentiles. Wilcoxon rank-sum test was used for statistical analysis. n.s. p>0.05, ****p<0.0001. **(C)** Volcano plot representing gene expression changes of upregulated genes in BCOR^KO^ vs. WT (left panel) and BCOR^KO^/BCORL1^KO^ vs. WT (right panel) K562 cells. Only PRC1.1 responsive genes of cluster C1 (n=65) are shown. Log2 fold changes of 1 and padj values of 0.05 are depicted by dotted lines. **(D)** ChIP-seq and ATAC-seq coverage data tracks of the PRC1.1 responsive gene *FGFR1* in WT or PRC1.1 mutant K562 cells. **(E)** *FGFR1* gene expression levels (normalized DESeq2 counts) in WT or PRC1.1 mutant K562 cells (upper panel) represented as mean ± SEM. Corresponding western blot analysis of FGFR1 protein levels (lower panel) (FGFR1: 91kDa, glycosylated FGFR1: 120kDa). **(F)** Schematic of the experimental workflow for *in vitro* competition assays. 80% RFP657 positive control cells were mixed with 20% BFP positive competitor cells and cultured in the presence of DMSO or indicated drugs, with or without FGF2 supplementation (10ng/mL). Fluorophore expression was tracked over time by flow-cytometry analysis. Competitor conditions expressed non-targeting control, or sgRNAs targeting *BCORL1*, *BCOR*, *BCOR* and *BCORL1* or *KDM2B*. **(G)** Heatmaps representing competitor percentages over the course of 10 days. Cells were cultured in the presence of DMSO (upper panel) or 1µM imatinib (lower panel) with or without FGF2 supplementation (10ng/mL). **(H)** Percentages of competitor cells expressing sgRNAs targeting *BCOR/BCORL1* (highlighted in red) or non-targeting control (highlighted in dark grey) over time. Cells were treated with 1µM imatinib alone (solid line) or with 1µM imatinib and 1µM FGFRi (PD173074) in combination (dotted line).

Based on their sensitivity to partial PRC1.1 inactivation, responsive genes clustered into two groups: cluster 1 (C1, n=65) and cluster 2 (C2, n=567) (Figure 5D). C1 genes showed significantly higher expression levels in single BCOR^KO^ and double BCOR^KO^/BCORL1^KO^ cells compared with C2 genes, and were significantly more highly expressed in double BCOR^KO^/BCORL1^KO^ cells than in single BCOR^KO^ cells (Figure 7B). PRC1.1 reduction in single BCOR mutant cells is thus sufficient to activate a subset of highly responsive PRC1.1 target genes whereas complete PRC1.1 loss in BCOR^KO^/BCORL1^KO^ cells results in activation of the full set of target genes.

Within the C1 gene cluster, the fibroblast growth factor receptor 1 (FGFR1) was among the most highly upregulated genes in partially as well as fully deficient PRC1.1 mutant cells (Figures 7C and S6B). At baseline, the *FGFR1* locus had a repressed chromatin state marked by low accessibility, enrichment of H2AK119ub and H3K27me3, and absent H3K27ac (Figure 7D). Accordingly, there was no *FGFR1* transcription and no detectable FGFR1 protein (Figure 7E). Partial PRC1.1 inactivation in BCOR^KO^ and BCOR^PUFD-Tr^ cells caused significantly increased FGFR1 expression without epigenetic reprogramming, whereas complete disruption of PRC1.1 in BCOR^KO^/BCORL1^KO^, BCOR^PUFD-Tr^/BCORL1^KO^, PCGF1^KO^, and KDM2B^KO^ caused loss of repressive histone marks and increased chromatin accessibility that correlated with a significant increase in FGFR1 gene expression and protein levels (Figures 7D and 7E). To confirm this observation in an independent AML cell line model, we generated MOLM13 cells with BCOR^KO^, BCORL1^KO^, BCOR^KO^/BCORL1^KO^ or KDM2B^KO^, and found that PRC1.1 inactivation caused similar activation of FGFR1 expression (Figure S6C). To determine whether the effect of PRC1.1 inactivation on FGFR1 expression was reversible and maintained dependency on intact BCOR/PRC1.1, we expressed a sgRNA-resistant full-length BCOR cDNA in BCOR^KO^/BCORL1^KO^ cells. Complementation of BCOR expression restored PRC1.1 complex assembly (Figure S2G) and decreased FGFR1 protein levels in PRC1.1 deficient cells (Figure S6D).

### PRC1.1 mutations drive aberrant FGFR1 activation which mediates resistance to targeted therapy

Upregulation of FGFR1 expression has been reported to be a mechanism of resistance to tyrosine kinase inhibition (Javidi-Sharifi et al., 2019; Traer et al., 2014, 2016), and *BCOR* mutations have been specifically linked to CML blast phase and resistance to BCR-ABL-targeted inhibitors (Branford et al., 2018). We therefore hypothesized that PRC1.1 disruption drives resistance to targeted BCR-ABL inhibition in K562 cells via derepression of FGFR1. To test this hypothesis, we assessed the relative growth of PRC1.1 deficient cells compared with control cells using *in vitro* competition assays. We mixed fluorochrome-labeled control cells (Cas9 plus non-targeting sgRNA) and competitor cells (Cas9 plus sgRNA targeting *BCOR*, *BCORL1*, *BCOR*/*BCORL1*, or *KDM2B*) at an 80:20 ratio, then used flow cytometry to measure their relative proportion over 10 days with or without the addition of inhibitors (Figure 7F). In baseline conditions (media with 10% serum, DMSO), PRC1.1 mutated cells did not have a selective growth advantage compared with control cells (Figure 7G). In contrast, when we blocked BCR-ABL signaling with imatinib (1µM), BCOR^KO^, BCOR^KO^/BCORL1^KO^ and KDM2B^KO^ cells all showed selective growth advantage compared with control cells, indicating increased relative resistance to imatinib (Figure 7G). Single BCOR^KO^ cells that showed less pronounced activation of signaling genes including FGFR1 (Figures 7C and 7E), were less resistant to imatinib than double BCOR^KO^/BCORL1^KO^ or KDM2B^KO^ cells, reflected by lower proportion of BFP positive cells at day 10 (BCOR^KO^: 27.5%; BCOR^KO^/BCORL1^KO^: 52.1%; KDM2B^KO^: 49.7%) (Figure 7G). In each competitor condition, we observed a concomitant increase in indel fraction at the sgRNA target sites, confirming correlation between fluorescent markers and gene targeting (Figure S6E).

To determine whether the effect of PRC1.1 inactivation on imatinib resistance was mediated by FGFR1 activity, we measured functional consequences of augmenting or inhibiting FGFR1 signaling.

Supplementation of media with FGF2 (10ng/mL) augmented the magnitude of selective advantage of BCOR^KO^, BCOR^KO^/BCORL1^KO^ and KDM2B^KO^ cells compared with control cells (Figure 7G) and was correlated with high FGFR1 protein levels at day 13 (Figure S6F). In contrast, concurrent treatment of cells with imatinib and the selective small molecule FGFR1 inhibitor PD173074 (10ng/ml) completely abrogated resistance of BCOR^KO^/BCORL1^KO^ cells compared with BCOR^KO^/BCORL1^KO^ cells treated with imatinib alone (Figure 7H). Treatment of K562 cells with PD173074 alone had no effect on cell growth or survival (Figure S6G). Together, these data indicate that genetic inactivation of PRC1.1 drives imatinib resistance via its effects on FGFR1.

## Discussion

Here, we paired genetic analysis of 6162 primary patient samples with functional studies in cell line and primary AML samples to define the role of PRC1.1 alterations in the pathogenesis of myeloid malignancies. We found that BCOR and BCORL1 are central adaptors that are required for PRC1.1 complex assembly and target gene repression. Leukemia-associated *BCOR* and *BCORL1* mutations cooperate to drive disease progression by selectively unlinking the enzymatic RING/PCGF1 core from the complex, thereby causing a quantitative reduction of enzymatically active PRC1.1 at chromatin and driving aberrant activation of oncogenic signaling programs. We demonstrate further that derepression of PRC1.1 target genes in BCOR/BCORL1-mutated leukemia cells mediates a functional resistance to kinase inhibitor treatment that is reversible with FGFR1 inhibition.

Oncogenic gene mutations that introduce a premature termination codon can operate by multiple mechanisms, including loss-of-function or gain-of-function depending on the protein product that is expressed (Lindeboom et al., 2019; Yang et al., 2018). Here, we show that pathogenic *BCOR* mutations have diverse consequences on BCOR protein expression but share a unifying mechanism for their epigenetic and oncogenic effects. Specifically, leukemia-associated *BCOR* mutations all cause loss of a stabilizing interaction between the C-terminal BCOR-PUFD domain and the PRC1.1-specific adaptor PCGF1, which results in separation of the enzymatic core of PRC1 from the chromatin-targeted PRC1.1 auxiliary subcomplex. This paradigm is exemplified by a subset of *BCOR* mutations in patients that cause stable expression of PUFD-truncated proteins by escaping nonsense mediated decay. We show in both patient samples and engineered cell line models that these PUFD-truncating BCOR mutations disrupt PCGF1 binding and stability and result in derepression of a core set of PRC1.1 targets. Further, we found that expression of PUFD-truncated BCOR was functionally equivalent to complete BCOR loss, indicating that isolated disruption of the BCOR-PCGF1 protein-protein interaction is sufficient to drive pathogenicity in myeloid malignancies. Notably, C-terminal truncated BCOR that lost PCGF1 binding was still able to interact with KDM2B, in line with a recent study showing that exogenously expressed BCOR lacking PUFD bound KDM2B but not PCGF1 (Wang et al., 2018). The formation of a stable KDM2B-BCOR subcomplex in the absence of PCGF1 binding might have broader implications in oncogenesis. For example, internal tandem duplications (ITDs) in BCOR PUFD domains in patients with clear cell sarcoma of the kidney (CCSK), CNS-PNET, and other cancers are predicted to disrupt PCGF1 interaction (Wong et al., 2020). We speculate that loss of PCGF1 might allow oncoproteins to hijack BCOR-ITD-KDM2B subcomplexes, resulting in gain or altered PRC1.1 function, providing a conceptual model by which PRC1.1 alterations may drive oncogenesis in a cancer specific context.

Our study provides a mechanistic basis for the higher frequency of *BCOR* mutations in leukemia compared with *BCORL1*. We found that BCOR and BCORL1 participate in functionally redundant and mutually exclusive PRC1.1 species, sharing key protein interaction partners and localizing to the same set of chromatin loci. The aggregate PRC1.1 activity in a cell thus seems to reflect a composite of BCOR-PRC1.1 and BCORL1-PRC1.1 complex activity. We found further that BCOR was significantly higher expressed than BCORL1 in normal and malignant hematopoiesis and that *BCOR* mutations did not result in a compensatory increase of BCORL1 gene expression levels or *vice versa*. Consequently, *BCOR* mutations caused a more pronounced quantitative reduction of global PRC1.1 levels than *BCORL1* mutations that correlated with transcriptional activation of a set of highly responsive PRC1.1 target genes. Our data indicate that *BCOR* and *BCORL1* mutations cooperate by progressive reduction of PRC1.1 levels and a dose-dependent derepression of PRC1.1 target genes. In line with this model, *BCORL1* mutations commonly arose in the context of a preexisting *BCOR* mutation in patients with myeloid malignancies which was associated with disease progression.

We found that *PCGF1* or *KDM2B* were rarely mutated in patients with AML, but that biallelic genetic inactivation of *PCGF1* or *KDM2B* in our AML cell line model mimicked the functional consequences of *BCOR/BCORL1* mutations. *BCOR* and *BCORL1* are located on the X chromosome and have been reported not to escape from X-inactivation in cancer (Grossmann et al., 2011), indicating that complete functional disruption of *BCOR* or *BCORL1* can be achieved with a single mutational event in individuals with either XX or XY genotype. In contrast, *PCGF1* (chromosome 2) and *KDM2B* (chromosome 12) are both autosomal genes and would require two mutational events to confer target gene derepression that would lead to a similar clonal advantage. Consistent with this observation, the single *PCGF1* mutation we identified in the AML cohort was present at high variant allele fraction, suggesting loss of heterozygosity.

Previous studies indicated that H2AK119ub deposition by non-canonical PRC1 complexes was critical for target gene repression and PRC2 recruitment (Blackledge et al., 2020; Fursova et al., 2019; Tamburri et al., 2020). This was consistent with our finding that PRC1.1 disruption resulted in loss of H2AK119ub and reduced PRC2 binding at target genes which correlated with transcriptional activation. While we observed widespread binding of PRC1.1 to repressed promoters, only a subset of CpG rich PRC1.1 bound genes were derepressed upon PRC1.1 disruption. Synergistic as well as unique functions of non-canonical PRC1 complexes have been previously reported suggesting that PRC1.3/5 or PRC1.6 complexes might compensate for PRC1.1 loss at shared target genes (Scelfo et al., 2019; Tamburri et al., 2020). A recent study showed that PRC1 complexes promoted active gene expression in mouse embryonic epidermal progenitors (Cohen et al., 2018). While we observed PRC1.1 binding to active gene promoters in leukemia, we did not detect changes in expression levels of active genes upon PRC1.1 disruption. We therefore favor a model proposed by Farcas et al. (Farcas et al., 2012) in which PRC1.1 may sample CpG rich gene promoters for susceptibility to Polycomb mediated silencing independent of their chromatin states, whereas additional factors are required to facilitate gene repression.

The identification of specific gene mutations or gene expression signatures may inform drug responses in patients with myeloid malignancies (Tyner et al., 2018). We found in both cell line models and primary AML samples that PRC1.1 deficiency was associated with activation of cell signaling pathways regulating stem cell pluripotency, consistent with a previously proposed role of PRC1.1 in regulating stem cell transcriptional programs (Kelly et al., 2019; Wang et al., 2018). We further demonstrated that epigenetic activation of the PRC1.1 target gene FGFR1 mediated imatinib resistance in PRC1.1 deficient K562 cell lines that could be overcome by FGFR1 inhibition. Our data indicate that PRC1.1 deficient cells were able to maintain cell signaling and survival in the absence of BCR-ABL signaling by utilizing FGFR1 as an alternative signaling pathway. This was supported by previous studies demonstrating that activation of FGFR signaling was a mechanism of TKI resistance in K562 cells (Traer et al., 2014) and that *BCOR* and *BCORL1* mutations were associated with blast crisis in CML patients (Branford et al., 2018). Our results further suggest that *BCOR* and *BCORL1* mutations may define more broadly a group of PRC1.1-deficient myeloid malignancies that respond to pharmacologic inhibition of hyperactive RAS/MAPK signaling.

## Acknowledgements

This work was supported by the National Institutes of Health [K08CA204734 (R.C.L.), R01HD084459 (V.J.B.)], the Edward P. Evans Foundation (R.C.L.), the German Research Foundation [DFG-SCHA2121/1-1 (E.J.S.)], the Research School for Translational Medicine Goettingen (E.J.S), the Dana-Farber Cancer Center Center for Cancer Genome Discovery, the Ted and Eileen Pasquarello Tissue Bank in Hematologic Malignancies, and the DFCI Hematologic Malignancies Data Repository.

## Author Contributions

Conceptualization, E.J.S and R.C.L.; Methodology, E.J.S., H.C.W., and I.F.; Formal Analysis, E.J.S., R.C.L., H.C.W., A.A., Y.X., and E.R.A.; Investigation, E.J.S., H.C.W., K.L., A.A., and E.R.A.; Resources, M.D.G., V.J.B., C.A.M., H.M.M., E.S.Wang, L.P.G., M.P.C., R.S.V., E.S.Winer, J.S.G., R.M.S., and M.R.L., Writing – Original Draft, E.J.S and R.C.L.; Writing – Review & Editing, E.J.S, R.C.L, H.C.W., C.A.M., P.C., M.D.G., E.R.A., I.F., C.J.G, M.S., E.S.Wang, L.P.G., M.P.C., R.S.V., E.S.Winer, J.S.G., R.M.S, M.R.L., S.A.C., H.W.L., V.J.B., and M.E.F.; Supervision, R.C.L., C.A.M., M.D.G., M.S., S.A.C, P.C., H.W.L, V.J.B., and M.E.F.; Funding Acquisition, E.J.S and R.C.L.

## Methods

### Lead Contact and Materials Availability

Further information and requests for resources and reagents should be directed to and will be fulfilled by the Lead Contact, Coleman Lindsley.

### Experimental Model and Subject Details Patient samples

We prepared DNA from bone marrow aspirate samples obtained from 433 patients with newly diagnosed AML prior to treatment. We performed targeted sequencing of 113 genes known to be recurrently mutated in AML or in germline syndromes predisposing to development of myeloid malignancies.

For RNA-seq, ChIP-seq, IP and western blot analysis, samples were obtained with informed consent from patients with newly diagnosed or relapsed acute myeloid leukemia. Leukemic blasts were isolated from whole blood using CD3 depletion (StemCell Technologies) and ficoll density gradient separation (GE Healthcare). Cell viability and sample purity were determined by trypan blue staining and FACS analyses. Cells were cryopreserved in FBS with 10% DMSO and stored in liquid nitrogen.

### Cell lines

K562 cells (female) were acquired from the Broad Institute, MV4-11 (male) and MOLM14 cells (male) from Dr. James Griffin’s lab. KG-1 (male) and OCI-AML3 cells (male) were purchased from ATCC. MOLM13 cells (male) were purchased from DSMZ. Cells were cultured in RPMI-1640 medium (Gibco^™^) with 10% fetal bovine serum (FBS, Sigma-Aldrich) and 1% penicillin-streptomycin-glutamine (PSG, Gibco^™^) at 37°C and 5% CO2. HEK293T cells were acquired from Dr. Benjamin Ebert’s lab and maintained in DMEM medium (Gibco^™^) with 10% FBS and 1% PSG at 37°C and 5% CO2.

## Method Details

### DNA sequencing and mutation analysis

Native genomic DNA was sheared and the library constructed per manufacturer protocol (Agilent). Libraries were then quantified and pooled up to 24 samples per lane in equimolar amounts totaling 500ng of DNA. Each pool was then hybridized to Agilent Custom SureSelect In Solution Hybrid Capture RNA baits. Each capture reaction was washed, amplified, and sequenced on an Illumina HiSeq 2000 100bp paired end run.

Fastq files were aligned to hg19 version of the human genome with BWA version 0.6.2 (Li and Durbin, 2009). Single nucleotide and small insertion and deletion calling was performed with samtools version 0.1.18 mpileup (Li et al., 2009) and Varscan version 2.2.3 (Koboldt et al., 2012). Pindel version 0.2.41,2 (Ye et al., 2009) was used for FLT3-ITD calling at the specific genomic locus located at chromosome 13:28,608,000-28,608,600. Variants were annotated to include information about cDNA and amino acid changes, sequence depth, number and percentage of reads supporting the variant allele, population allele frequency in the genome aggregation database (gnomAD) and presence in Catalogue of Somatic Mutations in Cancer (COSMIC), version 64. Variants were excluded if they had fewer than 15 total reads at the position, had fewer than 5 alternate reads, had variant allele fraction <2%, fell outside of the target coordinates, had excessive read strand bias, had excessive number of calls in the local region, caused synonymous changes, or were recurrent small insertions/deletions at low variant allele fraction adjacent to homopolymer repeat regions. No germline tissue was available for evaluation of somatic status of mutations.

### Generation of PRC1.1 mutant cell lines

PRC1.1 mutant cell lines were generated using CRISPR/Cas9 ribonucleoprotein transient transfection or stable lentiviral transduction of a sgRNA expressing construct.

For CRISPR/Cas9 ribonucleoprotein transient transfection, CRISPR RNA (crRNA) targeting BCOR, PCGF1, or control were mixed in equimolar concentrations with trans-activating CRISPR RNA (tracrRNA) and heated at 95°C for 5 minutes. Ribonucleoprotein (RNP) complexes were prepared by mixing complexed crRNA:tracrRNA oligos, Cas9-NLS protein, and Cas9 working buffer followed by incubation at room temperature for 20 minutes. K562 cells were resuspended in Nucleofector SF solution and then mixed with RNP complexes. The nucleofection mix was transferred to a Nucleocuvette® strip and run on a Nucelofector® 4D device.

For lentiviral production and stable cell line transduction, lentiviral vectors containing sgRNA targeting *BCOR*, *BCORL1*, *PCGF1*, *KDM2B*, or control were packaged using HEK293T cells. Lentiviral vectors were concentrated by ultracentrifugation at 25,000rpm for 2 hours, resuspended in plain DMEM (Gibco^™^), and stored at -80°C. For lentiviral infection, K562, MOLM13 or MOLM14 cells were plated with 4µg/ml polybrene (MilliporeSigma) and prepared lentivirus and spinfected at 2000rpm, 37°C for 2 hours. After 48-hour incubation, cells were selected based on antibiotic-resistance using 2 µg/ml puromycin (MilliporeSigma) or fluorophore expression by FACS sorting using Aria II SORP (BD).

PRC1.1 mutant K562 single-cell clones were derived from parental K562 cells harboring an N-terminal in-frame V5-tag at the endogenous *BCOR* locus, integrated by Cas9 RNP mediated Homologous Directed Repair (HDR) as previously described (Richardson et al., 2016). Single-cell clones were isolated by limiting dilution, expanded, and screened by Sanger sequencing and TIDE analysis (Brinkman et al., 2014). Single-cell clones with homozygous or compound heterozygous mutations in the respective genes were selected for further analysis.

### Exogenous BCOR expression

HA-tagged BCOR PUFD and CT domain cDNA were synthesized by Twist Bioscience. PUFD-, CT-, and 3xFLAG-tagged full-length BCOR cDNA were cloned into pXL303 plasmids and packaged into lentiviral vectors using HEK293T cells. Wild-type and BCOR/BCORL1 double knockout K562 cells were transduced as described above. Transduced cells were selected using 10 µg/ml blasticidin (Gibco^™^).

### *In-vitro* competition assay

Cas9 expressing K562 cells were transduced with a lentiviral vector expressing a BFP or RFP657 fluorophores and sgRNAs targeting *BCOR*, *BCORL1*, *BCOR* and *BCORL1*, *KDM2B*, or control. Cells expressing control or competitor sgRNAs were mixed in a 5:1 ratio and treated with either 0.01% DMSO, 1 µM imatinib (Cell Signaling Technology), 1 µM PD173074 (StemCell Technologies) or 1 µM imatinib/PD173074, with or without 10 ng/ml FGF2 (Cell Signaling Technology) supplementation. Every 48-72 hours, cells were analyzed by flow cytometry using LSRFortessa (BD) and cell cultures were replenished with fresh medium. Diva software was used for data acquisition and FlowJo v10 for data analysis.

### Cell viability analysis

2x10^4^ cells were seeded in 50 µl media per well in a 96-well flat-bottom plate. Media was supplemented with 10 ng/ml FGF2. DMSO or FGFR inhibitor (PD173074) was added in limiting dilutions to the wells. Cell viability was analyzed after 72h using CellTiter-Glo® Luminescence Cell Viability Assay (Promega). Cell viability was calculated relative to DMSO control.

### Co-Immunoprecipitation

For nuclear protein extraction, cells were washed with cold PBS and resuspended in cold Buffer A (10mM HEPES, 10mM KCl, 0.1mM EDTA, 0.1mM EGTA, 1x protease inhibitor cocktail (Thermo Scientific)). Cells were allowed to swell on ice for 15 min before 10% IGEPAL® (Sigma-Aldrich) was added. Lysates were pelleted at 14,000rpm, 4°C for 5 min. Nuclei were washed with cold PBS and dissolved in modified RIPA buffer (0.1% IGEPAL®, 0.1% Sodium deoxycholate, 150mM NaCL, 50mM Tris pH7.5, 1U/µl Benzonase® (MilliporeSigma), 1mM MgCl2, 2x protease inhibitor cocktail (Thermo Scientific)) at 4°C for 30 minutes with shaking. Cell debris was removed by centrifugation at 14,000 rpm, 4°C for 15 min. Protein concentrations in supernatants were quantified using the BCA assay kit (Thermo Fisher Scientific). 1 mg nuclear extracts, 3.6µg primary antibody or IgG control, and Dynabeads® Protein G (Invitrogen) were incubated overnight at 4°C with shaking. Beads were washed 3 times with wash buffer (150nM NaCl, 50mM Tris pH7.5) supplemented with 1% IGEPAL® and 3 times with wash buffer only. Beads were eluted with LDS sample buffer (Bio-Rad Laboratories) for western blot analysis or resuspended in wash buffer for subsequent mass spectrometry analyses. Antibodies used for IP: BCOR (Proteintech Cat#12107-1-AP, RRID:AB_2290335), BCORL1 (Bardwell Lab, RRID:AB_2889363), PCGF1 (Bardwell Lab, RRID: AB_2716801, (Gearhart et al., 2006)), KDM2B (Millipore Cat#09-864, RRID:AB_10806072), HA-tag (Abcam Cat#ab9110, RRID:AB_307019).

### Western blotting

For whole cell lysates, cells were counted and washed twice with cold PBS. Cell pellets were resuspended in SDS loading buffer (Bio-Rad Laboratories). Nuclear extracts were prepared as described above. Cell lysates were separated on 4%-12% SDS-PAGE gels and transferred to PVDF membranes. Membranes were blocked in 5% milk in TBST (1X Tris-Buffered Saline, 0.1% Tween® 20 Detergent) for 1 hour at room temperature and incubated with primary antibody overnight at 4°C. Membranes were then washed three times with TBST and incubated with respective secondary antibodies for 45 minutes at room temperature. Subsequently, membranes were washed three times with TBST and incubated with SuperSignal West Dura or Femto substrate (Thermo Fisher Scientific) for 30 seconds. Images were captured using Bio-Rad ChemiDoc MP. Membranes were stripped using Restore™ Plus Stripping buffer (Thermo Scientific) per manufacturer’s recommendations. ß-Actin or GAPDH protein levels were used as loading controls for whole cell lysates. Lamin B1 (LMNB1) was used as a loading control for nuclear extracts. Antibodies used for WB: BCOR (Proteintech Cat#12107-1-AP, RRID:AB_2290335), BCOR (Santa Cruz Cat#sc-514576, RRID:AB_2721913), BCOR (Bardwell Lab; Gearhart et al., 2006, RRID: AB_2716801), BCORL1 (Bardwell Lab, RRID:AB_2889363), PCGF1 (Santa Cruz Cat#sc-515371, RRID:AB_2721914), KDM2B (Millipore Cat#09-864, RRID:AB_10806072), RING1 (Abcam Cat#ab32644, AB_2238272), RING1B/RNF2 (Santa Cruz Cat#sc-101109, RRID:AB_1129072), H2AK119ub (Cell Signaling Technology Cat#8240, RRID:AB_10891618), H3K27me3 (Cell Signaling Technology Cat#9733,RRID:AB_2616029), BMI1 (Santa Cruz Cat#sc-390443), PCGF3 (Thermo Fisher Scientific Cat#PA5-96686, RRID:AB_2808488), PCGF5 (Abcam Cat#ab201511), RYBP (Sigma-Aldrich Cat#PRS2227, RRID:AB_1847589), CBX8 (Santa Cruz Cat#sc-374332, RRID:AB_10990104), SKP1 (Abcam Cat#ab76502, RRID:AB_1524396), HAUSP/USP7 (Abcam Cat#ab4080, RRID:AB_2214019), HA-tag (Abcam Cat#ab9110, RRID:AB_307019), FGFR1 (Cell Signaling Technology Cat#9740S, RRID:AB_11178519).

### Mass spectrometry

#### On-bead digest

Immunoprecipitation was performed using 1 mg input protein (nuclear extract) and 3.6 µg BCOR antibody (RRID:AB_2290335) or 3.6 µg IgG (RRID:AB_1031062) as described above. Rep1 and Rep2 samples were processed and analyzed independently on two consecutive days. Samples were lysed in modified RIPA buffer (0.1% IGEPAL®, 0.1% Sodium deoxycholate, 150mM NaCL, 50mM Tris pH7.5, 1U/µl Benzonase® (MilliporeSigma). Beads were washed 3 times with wash buffer (150nM NaCl, 50mM Tris pH7.5) supplemented with 1% IGEPAL® and 3 times with wash buffer only. The beads were resuspended in 20 μL of wash buffer, followed by 90 μL digestion buffer (2 M urea, 50 mM Tris HCl) and then 2 μg sequencing grade trypsin was added, followed by 1 hour of shaking at 700 rpm. The supernatant was removed and placed in a fresh tube. The beads were washed twice with 50 μl digestion buffer and combined with the supernatant. The combined supernatants were reduced (2 μl 500 mM dithiothreitol, 30 minutes, room temperature) and alkylated (4 μl 500 mM iodoacetamide, 45 minutes, dark), and a longer overnight digestion was performed: 2 μg (4 μl) trypsin, shaken overnight. The samples were then quenched with 20 μl 10% formic acid and desalted on 10 mg Oasis cartridges.

#### TMT labeling of peptides and basic reverse phase fractionation

Desalted peptides from each sample in each replicate were labeled with TMT reagents (Pierce/Thermo Fisher Scientific). Peptides were dissolved in 30 μl of 50 mM TEAB pH 8.5 solution (Sigma-Aldrich) and labeling reagent was added in 70 μl ethanol. After 1h incubation, the reaction was stopped with 50 mM Tris/HCl pH 7.5. Differentially labeled peptides were mixed and subsequently desalted on 10-mg Oasis cartridges according to the following protocol: cartridges were prepared for desalting by equilibrating with methanol, 50% ACN, 1% formic acid and 3 washes with 0.1% TFA. Desalted peptides were then labelled individually with TMT10 reagent according to the manufacturer’s instructions (Thermo Fisher Scientific).

For labeling with TMT, peptides from samples in each replicate were dissolved in 25 μl of 50 mM HEPES pH 8.5 and 0.2 mg of TMT labelling reagent was added to each sample in 10 μl of CAN as follows: channel 126: WT_IgG; channel 127N: WT_BCOR-IP; channel 127C: BCORKO_IgG; channel 128N: BCORKO_BCOR-IP; channel128C: PCGF1KO_IgG; channel 130N: PCGF1KO_BCOR-IP; channel 129C: BCORMUT_IgG (data not used in this manuscript); and channel 129N: BCORMUT_BCOR-IP (data not used in this manuscript). Samples were incubated with labelling reagent for 1 h with agitation. Next, the reaction was quenched with 2 μl of 5% hydroxylamine. Differentially labelled peptides were subsequently mixed, and the combined samples were fractionated into 6 fractions using basic reverse phase chromatography on Oasis Cartridges Samples were loaded onto the cartridge and washed 3 times with 1% formic acid. A pH switch was performed with 5 mM ammonium formate at pH 10, collected and run as fraction 1. Subsequent fractions were collected at the following ACN concentrations: 10% ACN in 5 mM ammonium formate; 20% ACN in 5 mM ammonium formate; 30% CAN in 5 mM ammonium formate; 40% ACN in 5 mM ammonium formate; and 50% ACN in 5 mM ammonium formate.

#### MS analysis

Reconstituted peptides were separated on an online nanoflow EASY-nLC 1000 UHPLC system (Thermo Fisher Scientific) and analyzed on a benchtop Orbitrap Q Exactive Plus mass spectrometer (Thermo Fisher Scientific). The peptide samples were injected onto a capillary column (Picofrit with 10 μm tip opening / 75 μm diameter, New Objective, PF360-75-10-N-5) packed in-house with 20 cm C18 silica material (1.9 μm ReproSil-Pur C18-AQ medium, Dr. Maisch GmbH, r119.aq). The UHPLC setup was connected with a custom-fit microadapting tee (360 μm, IDEX Health & Science, UH-753), and capillary columns were heated to 50°C in column heater sleeves (Phoenix-ST) to reduce backpressure during UHPLC separation. Injected peptides were separated at a flow rate of 200 nL/min with a linear 80 min gradient from 100% solvent A (3% acetonitrile, 0.1% formic acid) to 30% solvent B (90% acetonitrile, 0.1% formic acid), followed by a linear 6 min gradient from 30% solvent B to 90% solvent B. Each sample was run for 120 min, including sample loading and column equilibration times. The Q Exactive Plus instrument was operated in the data-dependent mode acquiring HCD MS/MS scans (R = 17,500) after each MS1 scan (R = 70,000) on the 12 most abundant ions using an MS1 ion target of 3 x 106 ions and an MS2 target of 5 x 104 ions. The maximum ion time utilized for the MS/MS scans was 120 msec; the HCD-normalized collision energy was set to 27; the dynamic exclusion time was set to 20 s, and the peptide match and isotope exclusion functions were enabled.

#### Quantification and identification of peptides and proteins

All mass spectra were processed using the Spectrum Mill software package v.6.01 pre-release (Agilent Technologies), which includes modules developed for iTRAQ and TMT6-based quantification. Precursor ion quantification was done using extracted ion chromatograms for each precursor ion. The peak area for the extracted ion chromatogram of each precursor ion subjected to MS/MS was calculated in the intervening high-resolution MS1 scans of the LC–MS/MS runs using narrow windows around each individual member of the isotope cluster. Peak widths in both time and m/z domains were dynamically determined on the basis of mass spectrometry scan resolution, precursor charge and m/z, subject to quality metrics on the relative distribution of the peaks in the isotope cluster versus theoretical. Similar MS/MS spectra acquired on the same precursor m/z in the same dissociation mode with ± 60 s were merged. MS/MS spectra with precursor charge >7 and poor quality MS/MS spectra, which failed the quality filter by having a sequence tag length less than 1, were excluded from searching.

For peptide identification, MS/MS spectra were searched against the human Uniprot database to which a set of common laboratory contaminant proteins was appended. Search parameters included: ESI-QEXACTIVE-HCD scoring parameters, trypsin or Lys-c/trypsin enzyme specificity with a maximum of 2 missed cleavage, 40% minimum matched peak intensity, ± 20 ppm precursor mass tolerance, ± 20 ppm product mass tolerance, and carbamidomethylation of cysteines and TMT-isobaric labelling of lysines and N-termini as fixed modifications. Oxidation of methionine, N-terminal acetylation and deamidated (N) were allowed as variable modifications, with a precursor MH+ shift range from −18 to 64 Da. Identities interpreted for individual spectra were automatically designated as valid by optimizing score and delta rank1– rank2 score thresholds separately for each precursor charge state in each LC–MS/MS run, while allowing a maximum target-decoy-based false-discovery rate (FDR) of 1.0% at the spectrum level.

In calculating scores at the protein level and reporting the identified proteins, redundancy is addressed in the following manner: the protein score is the sum of the scores of distinct peptides. A distinct peptide is the single highest scoring instance of a peptide detected through an MS/MS spectrum. MS/MS spectra for a particular peptide may have been recorded multiple times (that is, from different precursor charge states, isolated from adjacent BRP fractions or modified by oxidation of Met), but are still counted as a single distinct peptide. When a peptide sequence over eight residues long is contained in multiple protein entries in the sequence database, the proteins are grouped together and the highest scoring one and its accession number are reported. In some cases in which the protein sequences are grouped in this manner, there are distinct peptides that uniquely represent a lower scoring member of the group (isoforms or family members). Each of these instances spawns a subgroup, and multiple subgroups are reported and counted towards the total number of proteins identified. TMT ratios were obtained from the protein comparisons export table in Spectrum Mill. To obtain TMT protein ratios, the median was calculated over all of the distinct peptides assigned to a protein subgroup in each replicate. We required each protein to be detected with two or more unique peptides. To enable precise quantification, we limited our analysis to peptides that are uniquely assigned to a specific protein isoform or family member. For statistical analysis, we used the Limma package (Ritchie et al., 2015) in R (https://www.r-project.org/) to calculate multiple comparison adjusted P values using a moderated t-test.

### Proteomics Analysis and Visualization

Non-human proteins, proteins with less than two unique peptides, and proteins not present in the current HGNC database of protein coding genes (https://www.genenames.org/cgi-bin/statistics) were removed from further analyses. Ratios of intensities between channels were median normalized. Statistical analyses were performed using a two-sample moderated t test from the R package limma (Ritchie et al., 2015) to estimate p values for each protein and false discovery rate (FDR) corrections were applied to account for multiple hypothesis testing. Figures were generated using the R package ggplot2 version 3.2.1 (Wickham, 2016).

### RNA-seq

Total RNA was extracted according to the manufacturer’s recommendations using RNeasy Plus Mini Kit (Qiagen) or High Pure RNA Isolation Kit (Roche). Libraries were prepared using Roche Kapa mRNA HyperPrep strand specific sample preparation kits from 200 ng of purified total RNA according to the manufacturer’s protocol on a Beckman Coulter Biomek i7. The finished dsDNA libraries were quantified by Qubit fluorometer, Agilent TapeStation 2200, and RT-qPCR using the Kapa Biosystems library quantification kit according to manufacturer’s protocols. Uniquely dual indexed libraries were pooled in equimolar ratios and sequenced on an Illumina NextSeq 550 with single-end 75bp reads or a NovaSeq 6000 with 50bp read pairs by the Dana-Farber Cancer Institute Molecular Biology Core Facilities.

### ChIP-seq

For histone ChIP-seq analysis, cells were cross-linked for 10 min with 1% formaldehyde at room temperature on a shaker at 850rpm. Crosslinked nuclei were quenched with 0.125 M glycine for 5 minutes at room temperature and washed with PBS (containing protease inhibitor (Roche) and HDAC inhibitor Sodium Butyrate (NaBut)). For ChIP-seq analysis of BCOR, BCORL1, RNF2, KDM2B, and SUZ12, cells were first fixed with 2 mM DSG (Thermo Scientific) for 45 min at RT, followed by formaldehyde cross-linking as described above. After fixation, pellets were resuspended in 500 µl of 1% SDS (50 mM Tris-HCl pH 8, 10 mM EDTA) and sonicated in 1 ml AFA fiber millitubes for 25 minutes using a Covaris E220 instrument (setting: 75 peak incident power, 5% duty factor and 200 cycles per burst). Chromatin was immunoprecipitated using primary antibody and Dynabeads® Protein A/G (Invitrogen). ChIP-seq libraries were made using the ThruPLEX DNA-seq 48D Rubicon kit and purified. 75-bp single-end reads were sequenced on an Illumina NextSeq instrument. Antibodies used for ChIP-seq: BCOR (Proteintech Cat#12107-1-AP; RRID:AB_2290335), BCORL1 (Bardwell Lab, RRID:AB_2889363), KDM2B (Millipore Cat#17-10264, RRID:AB_11205420), RING1B/RNF2 (Cell Signaling Technology Cat#5694S, RRID:AB_10705604), SUZ12 (Abcam Cat#ab12073, RRID:AB_442939), H2AK119ub (Cell Signaling Technology Cat#8240, RRID:AB_10891618), H3K27me3 (Cell Signaling Technology Cat#9733, RRID:AB_2616029), H3K27ac (Diagenode Cat#C15410196, RRID:AB_2637079).

### Assay for Transposase-Accessible Chromatin Sequencing (ATAC-seq)

ATAC-seq experiments were performed using an adjusted version of the Omni-ATAC protocol (Corces et al., 2017). 100,000 cells were resuspended in 50 µl cold ATAC-resuspension buffer (RSB, 10 mM Tris-HCl pH 7.4, 10 mM NaCl and 3 mM MgCl2 in water) containing 0.1% NP40, 0.1% Tween-20, and 0.01% Digitonin and incubated on ice for 3 minutes. After lysis, 1 ml ATAC-RSB containing 0.1% Tween-20 was added to the mixture and centrifuged for 10 minutes at 1,500 RCF in a pre-chilled (4 °C) fixed-angle centrifuge. Supernatant was removed and nuclei were resuspended in 50 µl transposition mix (2.5µl transposase Tn5 (100 nM final), 25 µl 2x TD buffer, 16.5 µl 1x PBS, 0.5 µl 1% Digitonin, 0.5 µl 10% Tween-20 in water) (Corces et al., 2017). Transposition reactions were incubated at 37 °C for 30 minutes in a thermomixer. Libraries were made according to protocol described by Buenrostro et al. (Buenrostro et al., 2015). 35-bp paired-end reads were sequenced on an Illumina NextSeq 500 system.

### Multiplexed enhanced reduced representation bisulfite sequencing (mERRBS)

DNA from K562 cells (n=3) was extracted using the DNA Isolation Kit for Cells and Tissues from Roche (Cat#118147700011). Multiplexed enhanced reduced-representation bisulfite sequencing (mERRBS) was performed as previously described (Garrett-Bakelman et al., 2015) by the University of Michigan Epigenomics Core. Libraries were sequenced on a NovaSeq 6000 with 100bp single-end reads by the University of Miami Hussman Institute for Human Genomics.

### RNA-seq data analysis

Raw reads were mapped to the GRCh38 build of the human genome by STAR (Dobin et al., 2013). Gene counts were obtained using featureCounts (Liao et al., 2014). Normalization and differential gene expression analysis was performed using the R package DESeq2 version 1.24.0 (Love et al., 2014).

Gene expression levels were represented as boxplots using normalized DESeq2 counts. Genes with expression fold changes greater than 2 and adjusted p values less than 0.05 were considered differentially expressed. Gene set enrichment analysis (GSEA) was carried out using the gseKEGG function from clusterProfiler version 3.12.0 (padj cutoff value = 0.05, minimum gene set size = 70) (Yu et al., 2012). The g:GOSt function from g:Profiler version e99_eg46_p14_f0f4439 (Raudvere et al., 2019) was used to detect significantly enriched KEGG pathways of PRC1.1 responsive genes (n=632, Figure S5B). For unsupervised hierarchical clustering analysis and heatmap generation in Figure 5D, the count data was transformed using variance stabilizing transformation (VST) and analyzed using the R pheatmap package version 1.0.12 with default settings (default distance measure: Euclidean distance) (Kolde, 2015).

Raw RNA-seq counts from the BeatAML study were obtained from the authors via the Vizome portal (http://www.vizome.org/aml/, (Tyner et al., 2018)). Only AML samples with available whole-exome sequencing data were used for further analysis. AML samples with gene fusions, relapse samples as well as serial samples were excluded from the analysis (number of samples after filtering: n=238).

Normalization and differential gene expression analysis were performed using the R package DESeq2 version 1.24.0 (Love et al., 2014).

### ChIP-Seq and ATAC-seq data analysis

Quality control and preprocessing of raw ChIP-seq and ATAC-seq data was performed using the ChiLin pipeline (Qin et al., 2016). Reads were mapped to the hg19 human reference genome with BWA version 0.7.8 (Li and Durbin, 2009). Mapped reads were indexed and sorted using samtools version 1.9 (Li et al., 2009). Peaks were called with MACS2 version 2.1.1 (https://github.com/macs3-project/MACS; (Zhang et al., 2008)) at -q 0.01. The --SPMR option was used to generate bedgraph and bigwig files normalized to 1 million reads. Coverage tracks of bigwig files were visualized using the Integrative Genomics Viewer (IGV; (Robinson et al., 2011)). Peak annotations and genomic distributions of high-confidence peaks (-LOG10(qvalue) >10) were determined using the R package ChIPSeeker version 1.20.0 (Yu et al., 2015).

To determine ChIP or ATAC enrichment at promoter regions, the average signal score was calculated at TSS +/-5kb using multiBigwigSummary from deepTools version 3.0.0 (with BED file and – outRawCounts options; (Ramírez et al., 2016)). Bigwig files normalized per million of mapped reads were used as input files.

Gene promoters with BCOR signal scores >0.1 in K562 WT cells were defined as PRC1.1 bound (n=11,934) (Figure 4A). H3K27ac signal scores in K562 WT cells were used to subdivide PRC1.1 bound genes into active (H3K27ac >0.1) or repressed (H3K27ac <0.1) genes (Figure 5A). PRC1.1 unbound gene promoters (n=6639) with H3K27ac signal scores <0.1 were classified as inactive (n=6269). PRC1.1 unbound gene promoters with H3K27ac signal scores >0.1 (n=370) were not included in the analysis. We used the same approach to define active, repressed, and inactive chromatin states in BCOR^WT^ primary AML cells (Figure S4C). For correlation analysis of BCOR- and BCORL1-ChIP-seq datasets, average signal scores at TSS +/- 5kb were calculated using multiBigwigSummary and plotted as pairwise scatter plots (Figure 4B and Figure S3B).

For boxplot representations in Figure 5G, multiBigwigSummary signal scores at PRC1.1 responsive genes (n=632) were shown relative to the signal scores at PRC1.1 non-responsive genes (n=1799). Heatmaps and average density profiles of ChIP- or ATAC-signal enrichment at promoter regions were generated using computeMatrix (reference-point --referencePoint TSS -a 5000 -a 5000 -bs 50) followed by plotHeatmap or plotProfile from deepTools (Ramírez et al., 2016). Bigwig files normalized per million of mapped reads were used as input files. The option –regionsFileName was used to plot predefined regions. CpG rich promoters (n=10,851) and CpG poor promoters (n=7352) were defined based on CpG islands data from the UCSC Genome Browser version 2009-03-08 (Gardiner-Garden and Frommer, 1987).

### DNA methylation data analysis

Sequencing reads were aligned against a bisulfite-converted human genome (hg19) using Bowtie version 1.2.2 (Langmead et al., 2009), Bismark version 0.4.1 (Krueger and Andrews, 2011), and cutadapt (Martin, 2011) and the following code: /path/amp-errbs/ --prefix=myprefix --indir=/input_dir/ -- illumina=1.9 --adapter=NNAGATCGGAAGAGCACACGTCTGAACTCCAGTCAC --cutadapt -- genomePath=/path/hg19chromFa. The function ‘tiling window analysis’ from the R package methylKit version 1.10.0 (Akalin et al., 2012) was used to summarize methylation information within 200bp tiles. Methylation call files were then transformed to bed files for visualization using the Integrative Genomics Viewer (IGV; (Robinson et al., 2011)). The methylation status was represented with an 11-color gradient. Genomic coordinates, length and CpG counts of CpG islands were obtained from the CpG islands track from genome.ucsc.edu version 2009-03-08 (Gardiner-Garden and Frommer, 1987). Promoter regions (TSS +/-250bp) that overlapped with CGIs were determined with the BEDOPS closest-features function version 2.4.30 (Neph et al., 2012) and classified as CGI promoters (n=10,851).

### Quantification and Statistical Analysis

Data are represented as mean ± SD or mean ± SEM as indicated in the figure legends. Statistical parameters, including statistical significance and number of replicates, are described in the figure legends and in the method section. Statistical analyses were performed with GraphPad Prism software 8.4.3 (GraphPad Software, San Diego, CA).

---

**Figure S1.**
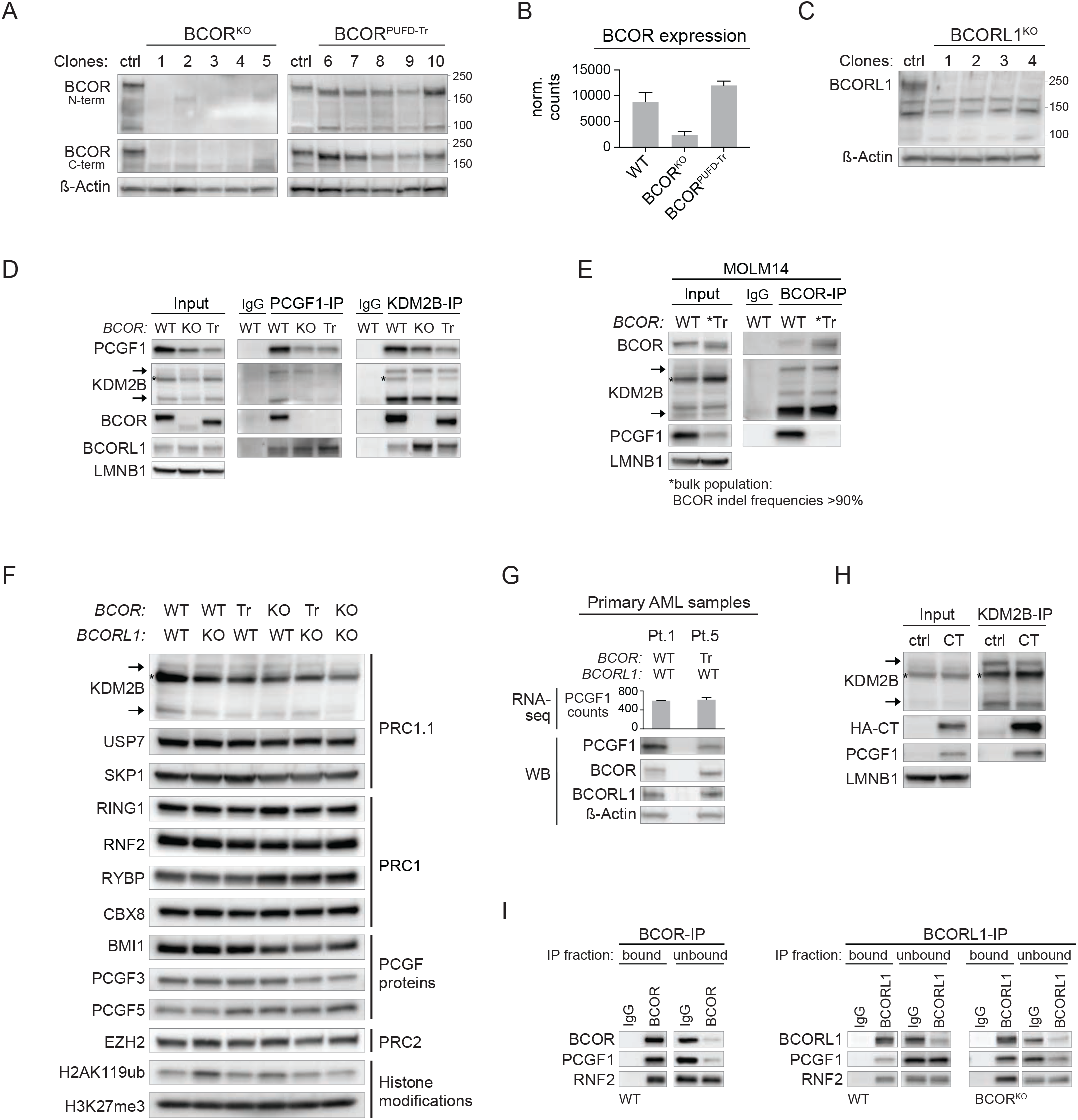
*BCOR* and *BCORL1* mutations disrupt assembly of distinct PRC1.1 complexes by disrupting binding to PCGF1. Related to Figure 2. **(A)** Western blot analysis of whole cell lysates from WT, BCOR^KO^ or BCOR^PUFD-Tr^ K562 single cell clones (related to Figure 2A). Antibodies recognizing N-terminal (amino acids 1-300, RRID:AB_2721913) or C-terminal (amino acids 1396-1755, RRID:AB_2290335) BCOR epitopes were used to detect truncated BCOR proteins. Predicted molecular weights: BCOR-WT: 1755aa/190kDa; BCOR^PUFD-Tr^: 1645aa/180kDa. **(B)** *BCOR* gene expression (DESeq2 normalized counts) in WT, BCOR^KO^ or BCOR^PUFD-Tr^ K562 cells represented as mean ± SD. **(C)** Western blot analysis of whole cell lysates from WT or BCORL1^KO^ K562 single cell clones (related to Figure 2A). Predicted molecular weight BCORL1-WT: 1785aa/190kDa. **(D)** PCGF1- and KDM2B-Co-IP-western blot analysis (nuclear extracts) in WT, BCOR^KO^ (KO), or BCOR^PUFD-Tr^ (Tr) K562 cells. KDM2B short (90kDa) and long (152kDa) isoforms are marked by arrows (asterisk indicates non-specific band). **(E)** BCOR-Co-IP-western blot analysis in WT or BCOR^PUFD-Tr^ (Tr) MOLM14 cells. BCOR^PUFD-Tr^ MOLM14 cells represent a bulk population with BCOR exon 14 indel frequencies of >90% determined by Sanger sequencing and TIDE analysis (Brinkman et al., 2014). KDM2B short (90kDa) and long (152kDa) isoforms are marked by arrows (asterisk indicates non-specific band). **(F)** Western blot analysis of nuclear extracts from WT, BCORL1^KO^, BCOR^PUFD-Tr^, BCOR^KO^, BCOR^PUFD-Tr^/BCORL1^KO^, or BCOR^KO^/BCORL1^KO^ K562 cell lines (related to Figure 2C; shared loading control: LMNB1). KDM2B short (90kDa) and long (152kDa) isoforms are marked by arrows (asterisk indicates non-specific band). **(G)** PCGF1 gene expression levels (DESeq2 normalized counts) in BCOR-WT (Pt.1) or BCOR^PUFD-Tr^ (Tr, Pt.5; BCOR p.L1647fs*3) AML patient cells represented as mean ± SD (upper panel). Lower panel shows corresponding western blot analysis of whole cell lysates. **(H)** KDM2B-Co-IP-western blot analysis in BCOR^KO^/BCORL1^KO^ K562 cells expressing HA-BCOR-CT (1462-1755aa). BCOR^KO^/BCORL1^KO^ cells transduced with a control vector were used as a negative control. **(I)** Western blot analysis of bound and unbound IP fractions of BCOR- or BCORL1-Co-IP analysis in WT or BCOR^KO^ K562 cells. IgG was used as negative control.

**Figure S2.**
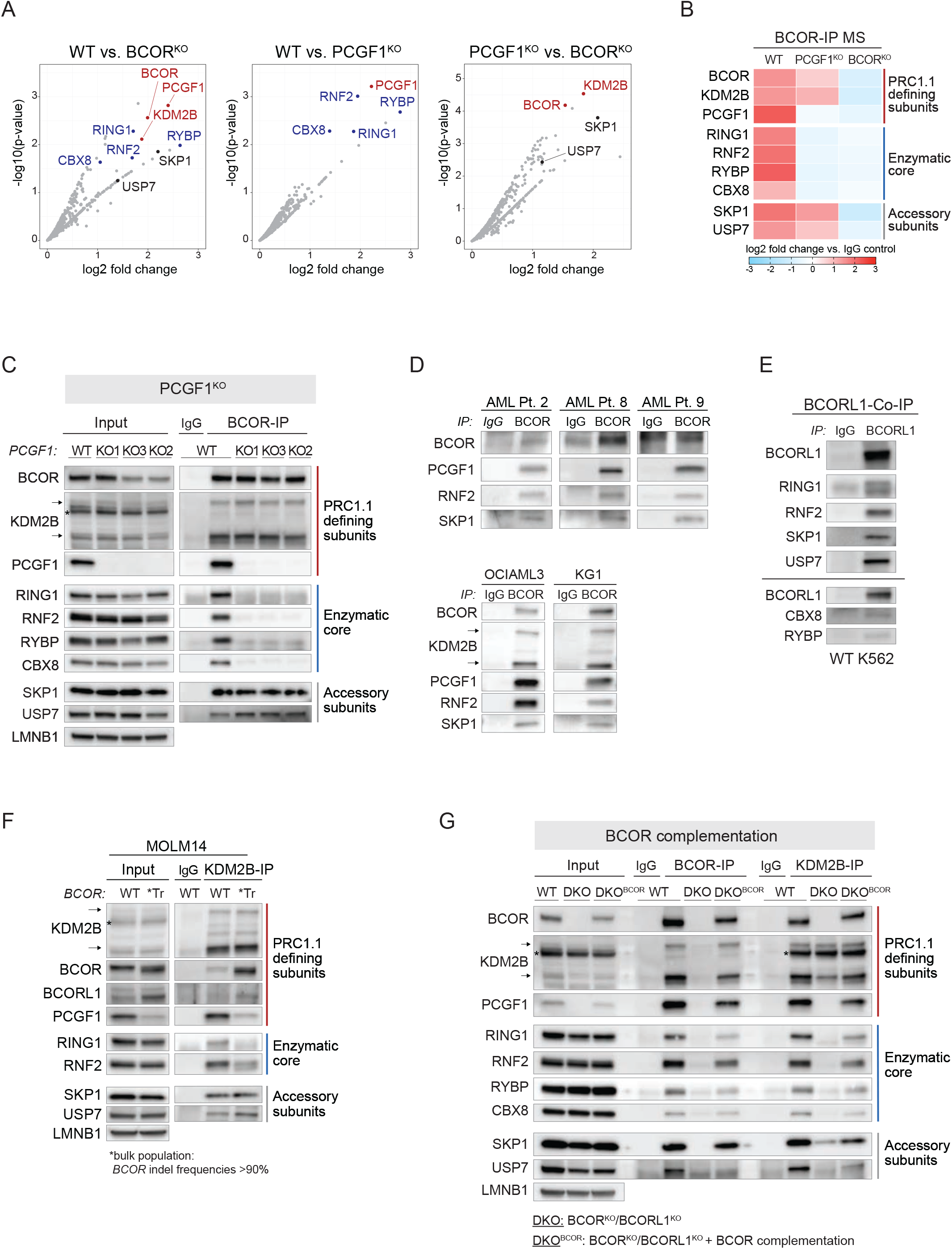
Loss of BCOR-PCGF1 or BCORL1-PCGF1 interaction disrupts complex assembly by uncoupling the enzymatic core from KDM2B. Related to Figure 3. **(A)** Enrichment of BCOR binding partners in BCOR-IP MS analysis. Volcano plots show -log10(p-value) against log2 fold change. n = 2 (process replicates for each condition). PRC1.1 defining subunits (BCOR, KDM2B, PCGF1) are highlighted in red, the enzymatic core (RING1, RNF2, RYBP, CBX8) in blue, and accessory subunits (SKP1, USP7) in black. **(B)** Enrichment of PRC1.1 subunits in BCOR-IP MS analysis in WT, PCGF1^KO^ or BCOR^KO^ K562 cells represented as log2 fold change compared to IgG control. n = 2 (process replicates for each condition). **(C)** BCOR-Co-IP-western blot analysis (nuclear extracts) in WT or PCGF1^KO^ K562 cells. The PCGF1 antibody RRID:AB_2721914 was used to detect PCGF1 protein levels (epitope: amino acids 128-247). KDM2B short (90kDa) and long (152kDa) isoforms are marked by arrows (asterisk indicates non-specific band). **(D)** Elution fractions of BCOR-Co-IP-western blot analysis in BCOR^WT^ primary AML patient cells (upper panel), or OCI-AML3 and KG1 AML cell lines (lower panel). KDM2B short (90kDa) and long (152kDa) isoforms are marked by arrows. **(E)** Elution fractions of BCORL1-Co-IP-western blot analysis in WT K562 cells. Black line indicates results from two independent experiments. **(F)** KDM2B-Co-IP-western blot analysis in WT or BCOR^PUFD-Tr^ (Tr) MOLM14 cells. BCOR^PUFD-Tr^ MOLM14 cells represent a bulk population with BCOR exon 14 indel frequencies of >90% determined by Sanger sequencing and TIDE analysis (Brinkman et al., 2014). KDM2B short (90kDa) and long (152kDa) isoforms are marked by arrows (asterisk indicates non-specific band). **(G)** BCOR- and KDM2B-Co-IP-western blot analysis in WT, BCOR^KO^/BCORL1^KO^ (DKO) or BCOR^KO^/BCORL1^KO^ cells complemented with WT-BCOR (DKO^BCOR^). Exogenous full-length WT-BCOR (BCOR isoform 1) was stably expressed in DKO^BCOR^ cells by lentiviral transduction.

**Figure S3.**
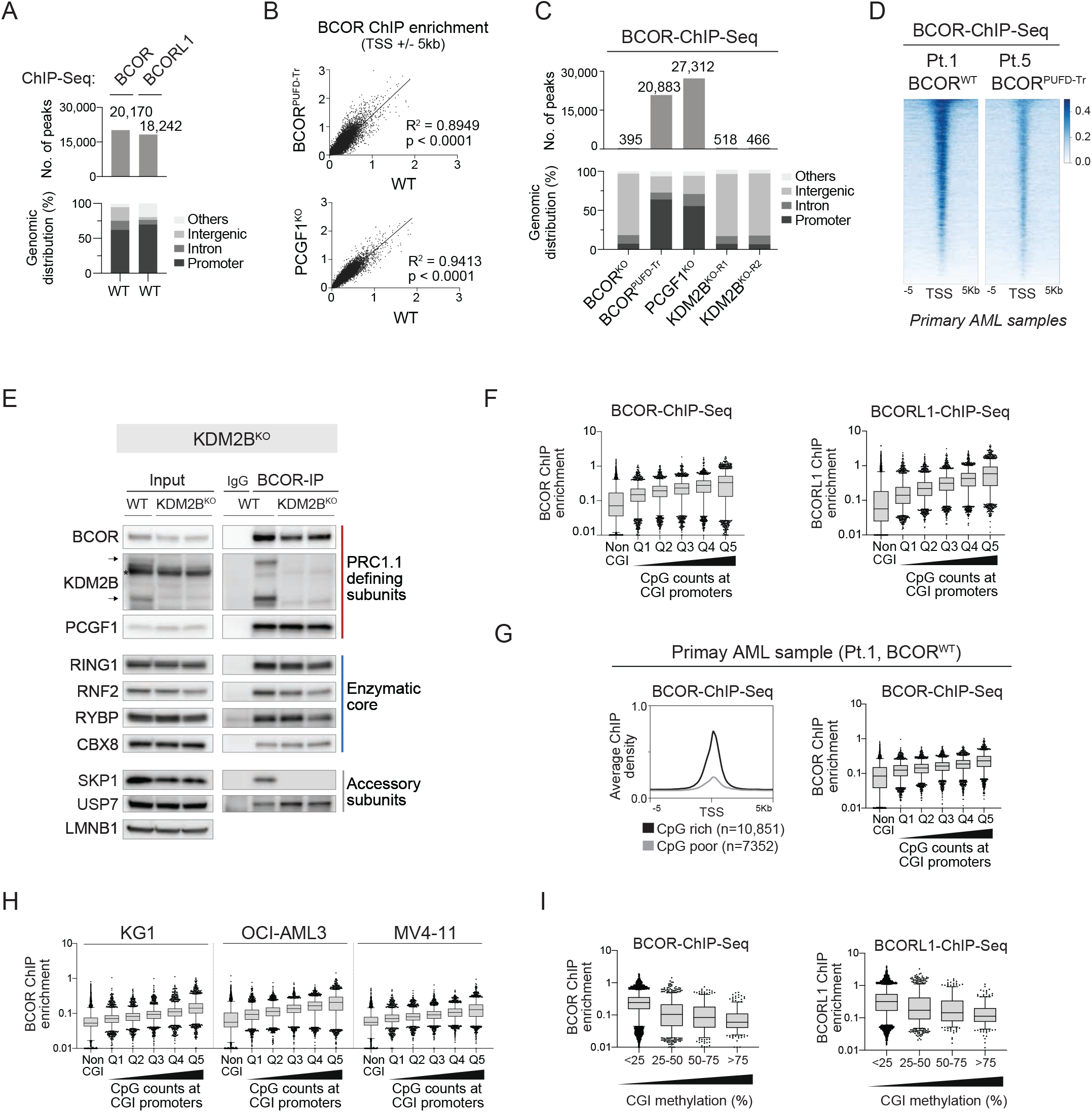
KDM2B is essential to recruit BCOR-PRC1.1 and BCORL1-PRC1.1 to unmethylated CpG island promoters. Related to Figure 4. **(A)** Total number of BCOR and BCORL1 peaks (threshold: -log10(q-value) >10) in WT K562 cells (upper panel). Distribution of BCOR and BCORL1 peaks in WT K562 cells according to their genomic location (lower panel). **(B)** Scatterplots showing the relationship between BCOR-ChIP-seq signals at TSS +/-5kb in BCOR^PUFD-Tr^ vs. WT K562 cells (upper panel), and in PCGF1^KO^ vs. WT K562 cells (lower panel). Pearson correlation coefficients and p values (two-tailed) as indicated. **(C)** Total number of BCOR peaks (threshold: -log10(q-value) >10) in BCOR^KO^, BCOR^PUFD-Tr^, PCGF1^KO^, or KDM2B^KO^ K562 cells (upper panel). Distribution of BCOR peaks according to their genomic location (lower panel). **(D)** Heatmap represents BCOR-ChIP enrichment in BCOR^WT^ (Pt.1) or BCOR^PUFD-Tr^ (Pt.5; BCOR p.L1647fs*3) primary AML patient samples at TSS +/-5kb. **(E)** BCOR-Co-IP-western blot analysis (nuclear extracts) in WT or KDM2B^KO^ K562 cells. KDM2B short (90kDa) and long (152kDa) isoforms are marked by arrows (asterisk indicates non-specific band). **(F)** Average BCOR ChIP-seq signals (left panel) and BCORL1 ChIP-seq signals (right panel) in WT K562 cells at non-CGI promoters (n=7414) or CGI promoters divided into quantiles based on CpG counts per CGI (Q1-4 n=2200; Q5 n=2199). CGI: CpG island. **(G)** Average density profiles of BCOR-ChIP signals in BCOR^WT^ (Pt.1) primary AML patient samples at CpG rich (n=10,851) or CpG poor (n=7352) promoters (left panel). Average BCOR ChIP-seq signals in BCOR^WT^ primary AML patient samples at non-CGI promoters (n=7414) or CGI promoters divided into quantiles based on CpG counts per CGI (Q1-4 n=2200; Q5 n=2199). **(H)** Average BCOR ChIP-seq signals in KG1, OCI-AML3, or MV4-11 AML cell lines at non-CGI promoters (n=7414) or CGI promoters divided into quantiles based on CpG counts per CGI (Q1-4 n=2200; Q5 n=2199). **(I)** Average BCOR- and BCORL1-ChIP-seq signals at CGI promoters subdivided based on average methylation percentages per CGI.

**Figure S4.**
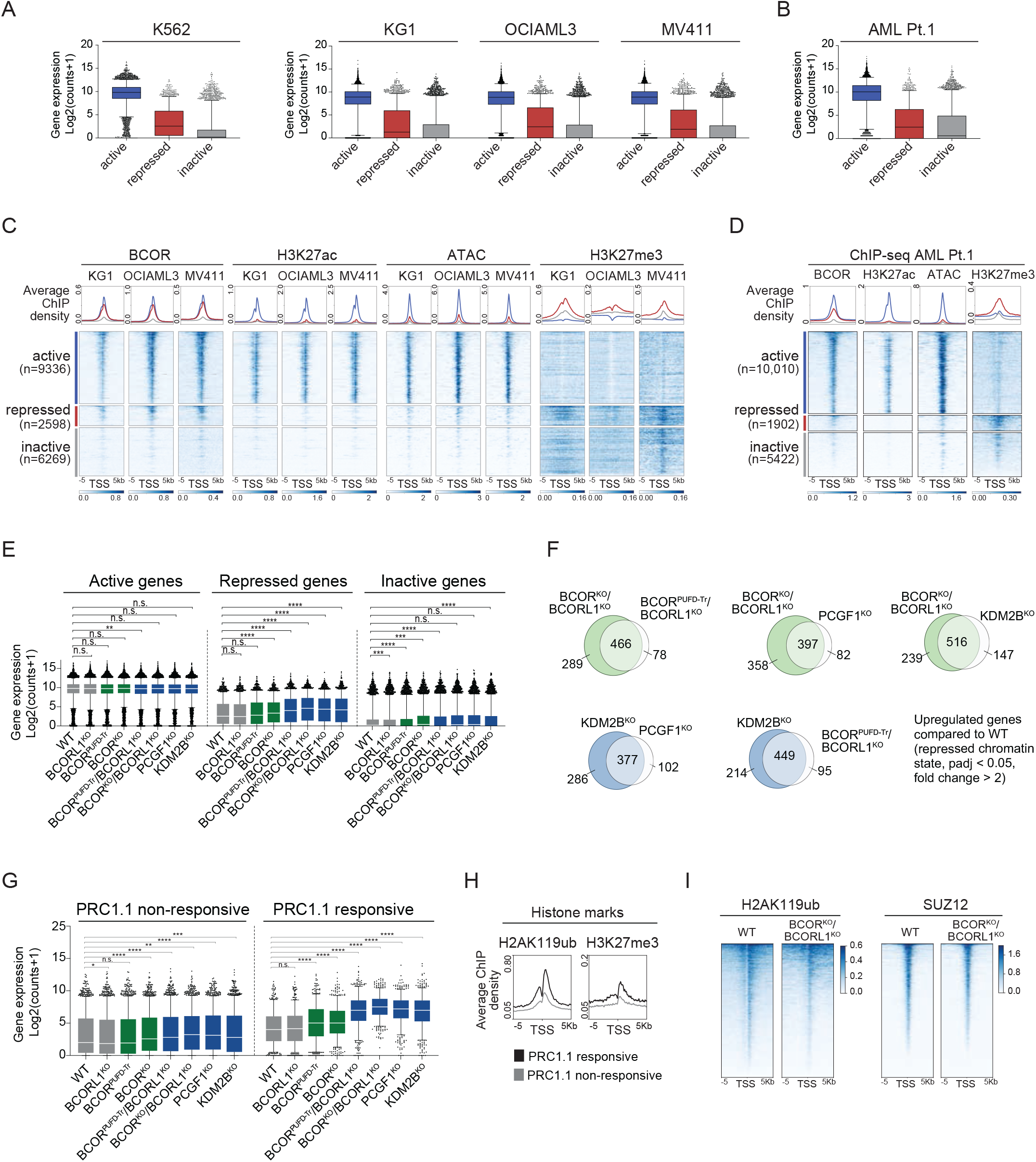
Recruitment of the PRC1.1 enzymatic core is required for target gene repression. Related to Figure 5. **(A)** Gene expression levels (DESeq2 normalized counts) of active, repressed or inactive genes in K562, KG1, OCI-AML3,or MV411 cell lines. BCOR- and H3K27ac-ChIP signal scores in K562 WT cells were used to define active, repressed, and inactive chromatin states (as depicted in Figures 5A and S4C). The line in the middle of the box represents the median, box edges represent the 25th and 75th percentiles, and whiskers show 5th and 95th percentiles. **(B)** Gene expression levels (DESeq2 normalized counts) of active, repressed or inactive genes in BCOR^WT^ primary AML cells (Pt.1). BCOR- and H3K27ac-ChIP signal scores were used to define active, repressed, and inactive chromatin states (as shown in Figure S4D). The line in the middle of the box represents the median, box edges represent the 25th and 75th percentiles, and whiskers show 5th and 95th percentiles. **(C)** Heatmaps and profiles of ChIP-seq signals of BCOR, histone modifications, and chromatin accessibility signals (ATAC) in KG-1, OCI-AML3, and MV4-11 cells at active, repressed, or inactive gene promoters (TSS +/-5kb). **(D)** Heatmaps and profiles of ChIP-seq signals of BCOR, histone modifications, and chromatin accessibility signals (ATAC) in BCOR^WT^ primary AML cells (Pt.1) at active, repressed, or inactive gene promoters (TSS +/-5kb). **(E)** Gene expression levels (log2 transformed DESeq2 normalized counts) of active (n=9335), repressed (n=2597) or inactive (n=6268) genes (as defined in Figure 5A) in WT or PRC1.1 mutant K562 cells. The line in the middle of the box represents the median, box edges represent the 25th and 75th percentiles, and whiskers show 5th and 95th percentiles. Wilcoxon rank-sum test was used for statistical analysis. n.s. p>0.05, *p<0.05, **p<0.01, ***p<0.001, ****p<0.0001. **(F)** Venn diagrams represent the overlaps of upregulated genes in PRC1.1 mutant cells vs. WT control (padj <0.05, fold change >2). Only genes that were in a repressed chromatin state at baseline were included in this analysis. **(G)** Box and whisker plots representing expression levels (log2 transformed DESseq2 normalized counts) of PRC1.1 non-responsive (n=1799) and PRC1.1 responsive genes (n=632) in WT or PRC1.1 mutant K562 cells (related to Figure 5D). The line in the middle of the box represents the median, box edges represent the 25th and 75th percentiles, and whiskers show 5th and 95th percentiles. Wilcoxon rank-sum test was used for statistical analysis. n.s. p>0.05, *p<0.05, **p<0.01, ***p<0.001, ****p<0.0001. **(H)** Average density profiles of histone ChIP-seq signals in WT K562 cells at PRC1.1 responsive (n=632) or PRC1.1 non-responsive genes (n=1799). **(I)** Heatmaps representing H2AK119ub- and SUZ12-ChIP signals at TSS +/-5kb in WT or BCOR^KO^/BCORL1^KO^ cells.

**Figure S5.**
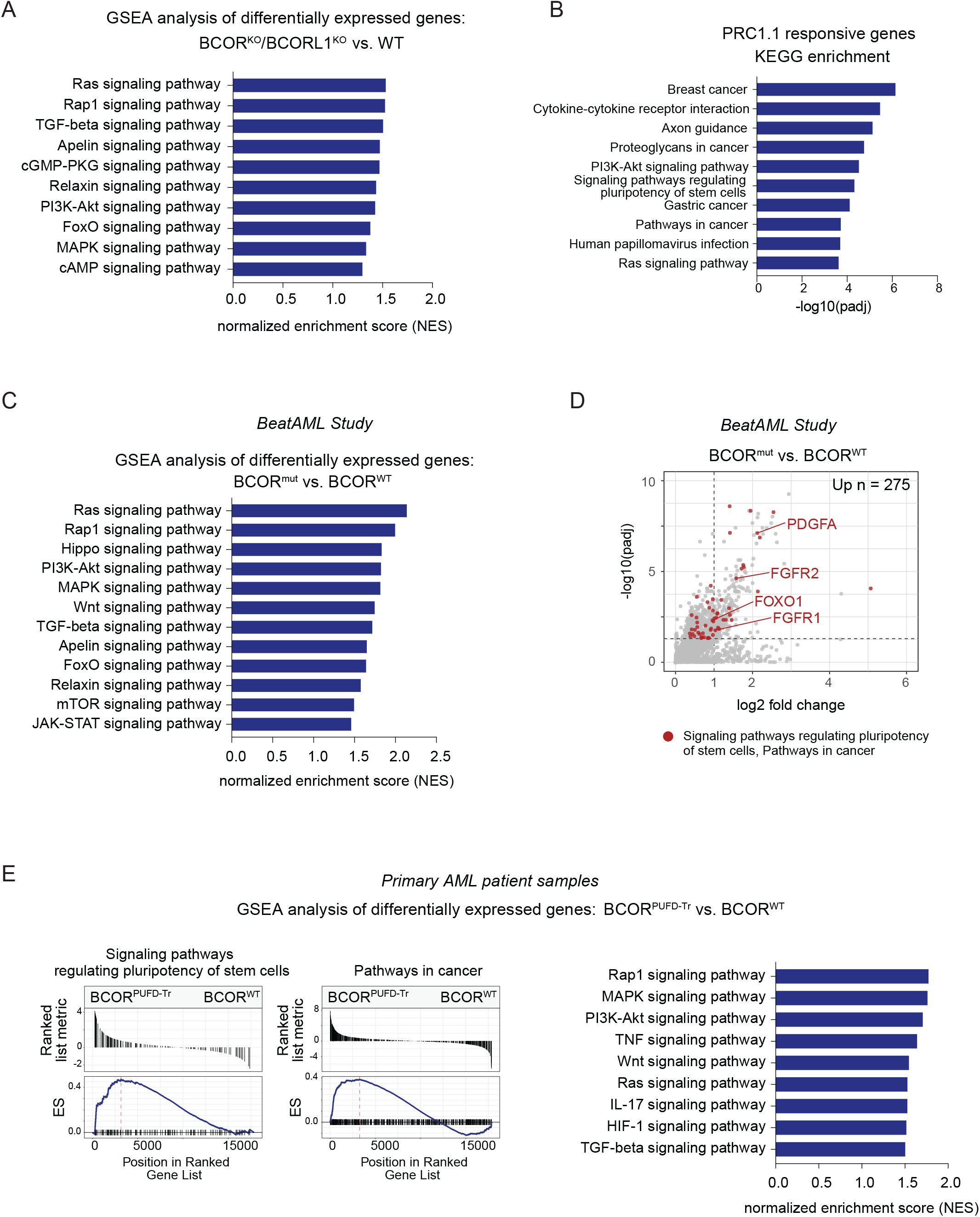
PRC1.1 regulates cell signaling programs in human leukemia. Related to Figure 6. **(A)** Overrepresented KEGG signaling pathways based on gene set enrichment analysis of differentially expressed genes in BCOR^KO^/BCORL1^KO^ compared to WT K562 cells. NES: normalized enrichment score. **(B)** KEGG pathway enrichment analysis of PRC1.1 responsive genes (n=632) using gProfiler (g:GOSt) (Raudvere et al., 2019). **(C)** Overrepresented KEGG signaling pathways based on gene set enrichment analysis of differentially expressed genes in BCOR^mut^ (n=22) compared to BCOR^WT^ (n=216) primary AML samples from the beatAML study (Tyner et al., 2018). NES: normalized enrichment score. **(D)** Volcano plot representing gene expression changes of upregulated genes in BCOR^mut^ (n=22) compared to BCOR^WT^ (n=216) primary AML samples from the beatAML study (Tyner et al., 2018). Genes represented in KEGG Signaling pathways regulating pluripotency of stem cells (hsa04550) or KEGG Pathways in cancer (hsa05200) are highlighted in red. Log2 fold changes of 1 and padj values of 0.05 are depicted by dotted lines. 275 genes were significantly upregulated more than 2-fold in BCOR^mut^ compared BCOR^WT^ cells. **(E)** KEGG gene set enrichment analysis of differentially expressed genes in BCOR^PUFD-Tr^ compared to BCOR^WT^ primary AML samples. The top plot represents the magnitude of log2 fold changes for each gene. The bottom plot indicates the enrichment score (ES) (left panel). Summary of overrepresented KEGG signaling pathways based on gene set enrichment analysis of differentially expressed genes in BCOR^PUFD-Tr^ compared to BCOR^WT^ primary AML samples (right panel). NES: normalized enrichment score.

**Figure S6.**
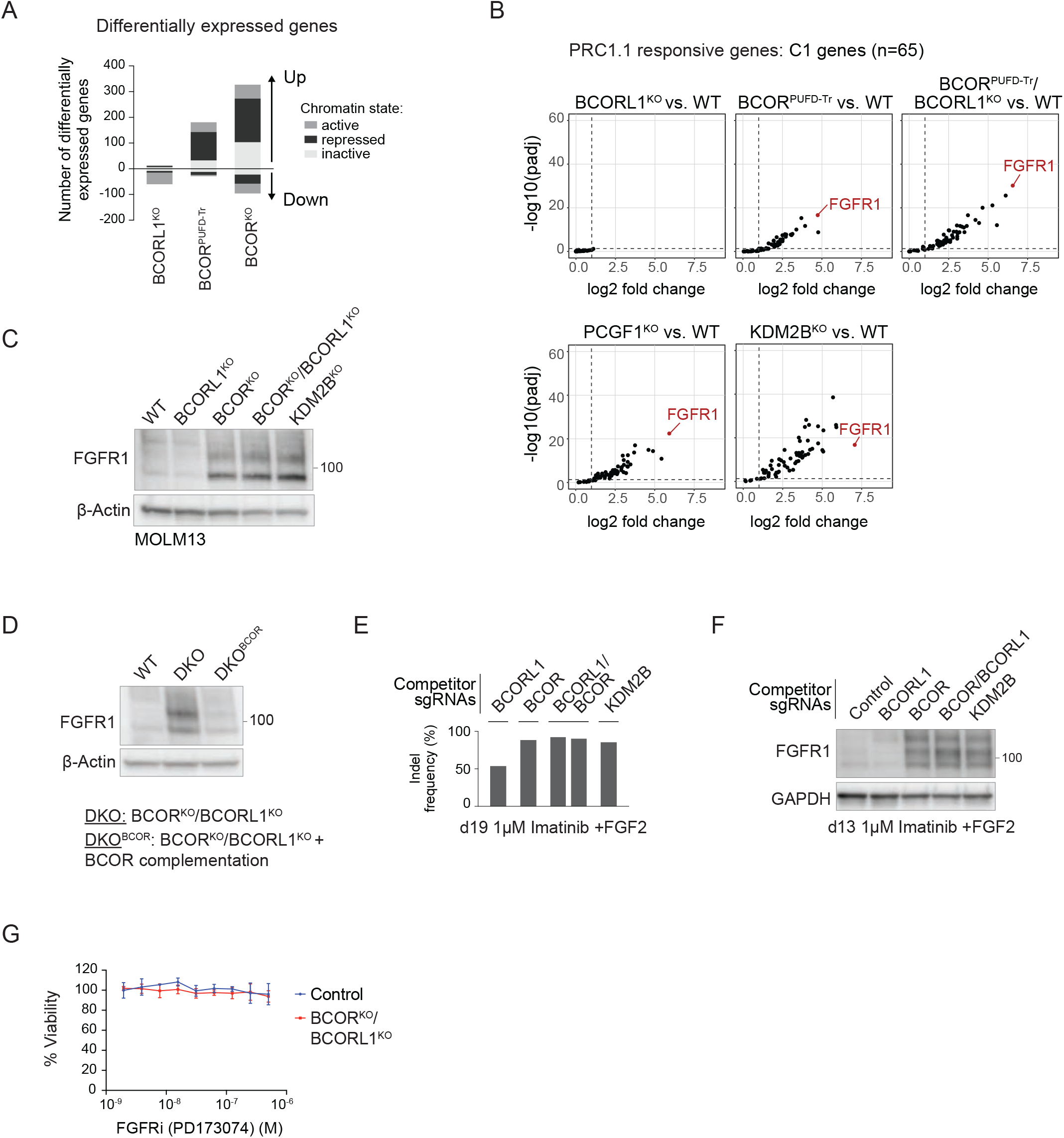
PRC1.1 mutations drive aberrant FGFR1 activation which mediates resistance to targeted therapy. Related to Figure 7. **(A)** Number of differentially expressed genes in PRC1.1 mutant compared to WT K562 cells (fold change >2, padj <0.05). Differentially expressed genes were subdivided based on their chromatin states defined in Figure 5A. **(B)** Volcano plots representing gene expression changes of upregulated genes in PRC1.1 mutant vs. WT K562 cells. Only PRC1.1 responsive genes of cluster C1 (n=65) are shown. Log2 fold changes of 1 and padj values of 0.05 are depicted by dotted lines. **(C)** Western blot analysis of FGFR1 protein levels in WT, BCORL1^KO^, BCOR^KO^, BCOR^KO^/BCORL1^KO^, or KDM2B^KO^ MOLM13 AML cell lines (FGFR1: 91kDa, glycosylated FGFR1: 120kDa). PRC1.1 mutant MOLM13 cells represent bulk populations. High indel frequencies (>90%) in respective genes were confirmed by Sanger sequencing and TIDE analysis (Brinkman et al., 2014). **(D)** Western blot analysis of FGFR1 protein levels in WT, BCOR^KO^/BCORL1^KO^ (DKO) or BCOR^KO^/BCORL1^KO^ K562 cells complemented with WT-BCOR (DKO^BCOR^). Exogenous full-length WT-BCOR was stably expressed in DKO^BCOR^ cells by lentiviral transduction. **(E)** Indel frequencies 19 days after *in vitro* competition of control versus competitor cells in the presence of 1µM imatinib and FGF2 ligand (10ng/mL) determined by Sanger sequencing and TIDE analysis (Brinkman et al., 2014). **(F)** Western blot analysis of FGFR1 protein levels 13 days after *in vitro* competition of control versus competitor cells in the presence of 1µM imatinib and FGF2 ligand (10ng/mL). FGFR1: 91kDa, glycosylated FGFR1: 120kDa, 140kDa. **(G)** Control or BCOR^KO^/BCORL1^KO^ cells were treated with DMSO or the FGFR inhibitor PD173074 at increasing concentrations for 72 hours. Cell viability was measured by using CellTiter-Glo luminescent assay (replicates n=2, error bars indicate standard deviation (SD)).

**Supplemental Table S1.** Related to Figure 1A. List of variants in genes encoding PRC1.1 subunits.

| Pat_ID | Gene | Chr | Start_pos | End_pos | Ref | Alt | cDNA change | AA change | Variant_Type | Reads_Ref | Reads_Alt | VAF |
| --- | --- | --- | --- | --- | --- | --- | --- | --- | --- | --- | --- | --- |
| AML_01 | BCOR | X | 39932868 | 39932868 | A | - | c.1731delT | p.N577fs | frameshift-del | 692 | 472 | 0.40 |
| AML_02 | BCOR | X | 39914767 | 39914767 | C | T | c.4494-1G>A | . | splice_site | 48 | 452 | 0.90 |
| AML_04 | BCOR | X | 39911550 | 39911550 | C | A | c.G5080T | p.E1694X | nonsense | 4 | 250 | 0.98 |
| AML_05 | BCOR | X | 39911610 | 39911610 | T | A | c.A5020T | p.K1674X | nonsense | 51 | 157 | 0.75 |
| AML_06 | BCOR | X | 39914707 | 39914708 | O | - | c.4654_4655del | p.Y1552fs | frameshift-del | 161 | 142 | 0.47 |
| AML_07 | BCOR | X | 39923016 | 39923016 | C | - | c.3692delG | p.R1231fs | frameshift-del | 442 | 460 | 0.51 |
| AML_08 | BCOR | X | 39933982 | 39933982 | - | CCCGGACC | c.616_617insGGTCCGGG | p.F206fs | frameshift-ins | 327 | 198 | 0.37 |
| AML_09 | BCOR | X | 39914625 | 39914625 | - | C | c.4736_4737insG | p.L1579fs | frameshift-ins | 25 | 273 | 0.92 |
| AML_10 | BCOR | X | 39932595 | 39932595 | - | AAGAC | c.2003_2004insGTCTT | p.V668fs | frameshift-ins | 681 | 207 | 0.23 |
| AML_11 | BCOR | X | 39931842 | 39931849 | O | - | c.2750_2757del | p.T917fs | frameshift-del | 249 | 223 | 0.47 |
| AML_12 | BCOR | X | 39913252 | 39913252 | T | - | c.4863delA | p.P1621fs | frameshift-del | 216 | 236 | 0.52 |
| AML_13 | BCOR | X | 39921556 | 39921556 | - | AC | c.4263_4264insGT | p.L1422fs | frameshift-ins | 688 | 13 | 0.02 |
| AML_13 | BCOR | X | 39921557 | 39921557 | - | GT | c.4262_4263insAC | p.R1421fs | frameshift-ins | 687 | 13 | 0.02 |
| AML_14 | BCOR | X | 39913588 | 39913588 | T | C | c.4640-2A>G | . | splice_site | 69 | 33 | 0.32 |
| AML_16 | BCOR | X | 39931996 | 39931996 | G | - | c.2603delC | p.T868fs | frameshift-del | 80 | 606 | 0.88 |
| AML_17 | BCOR | X | 39921391 | 39921391 | C | T | c.4326+1G>A | . | splice_site | 28 | 208 | 0.88 |
| AML_18 | BCOR | X | 39933205 | 39933205 | A | - | c.1394delT | p.V465fs | frameshift-del | 154 | 245 | 0.61 |
| AML_19 | BCOR | X | 39922123 | 39922123 | - | A | c.4048dupT | p.Y1350fs | frameshift-ins | 612 | 247 | 0.29 |
| AML_20 | BCOR | X | 39913528 | 39913528 | - | CGGGC | c.4799_4800insGCCCG | p.F1600fs | frameshift-ins | 512 | 101 | 0.16 |
| AML_21 | BCOR | X | 39931993 | 39931994 | O | - | c.2605_2606del | p.Y869fs | frameshift-del | 929 | 80 | 0.08 |
| AML_23 | BCOR | X | 39932002 | 39932002 | T | - | c.2597delA | p.H866fs | frameshift-del | 524 | 31 | 0.06 |
| AML_23 | BCOR | X | 39933541 | 39933541 | - | T | c.1057dupA | p.T353fs | frameshift-ins | 365 | 117 | 0.24 |
| AML_24 | BCOR | X | 39931848 | 39931848 | - | GA | c.2750_2751insTC | p.T917fs | frameshift-ins | 121 | 455 | 0.79 |
| AML_26 | BCOR | X | 39913252 | 39913252 | T | - | c.4863delA | p.P1621fs | frameshift-del | 45 | 397 | 0.90 |
| AML_27 | BCOR | X | 39933499 | 39933499 | G | - | c.1100delC | p.S367fs | frameshift-del | 174 | 291 | 0.63 |
| AML_27 | BCOR | X | 39933505 | 39933505 | - | TA | c.1093_1094insTA | p.S365fs | frameshift-ins | 119 | 291 | 0.67 |
| AML_29 | BCOR | X | 39913528 | 39913528 | - | A | c.4799dupT | p.F1600fs | frameshift-ins | 438 | 185 | 0.30 |
| AML_31 | BCOR | X | 39911489 | 39911489 | - | C | c.5140dupG | p.E1714fs | frameshift-ins | 322 | 38 | 0.11 |
| AML_31 | BCOR | X | 39914723 | 39914723 | G | A | c.C4639T | p.R1547X | nonsense | 352 | 24 | 0.06 |
| AML_31 | BCOR | X | 39916531 | 39916531 | - | G | c.4471_4472insC | p.N1491fs | frameshift-ins | 354 | 83 | 0.19 |
| AML_31 | BCOR | X | 39933301 | 39933301 | - | TGGCC | c.1297_1298insGGCCA | p.T433fs | frameshift-ins | 447 | 41 | 0.08 |
| AML_32 | BCOR | X | 39921391 | 39921391 | C | T | c.4326+1G>A | . | splice_site | 27 | 290 | 0.91 |
| AML_15 | BCORL1 | X | 129185992 | 129185992 | G | A | c.4853+1G>A | . | splice_site | 783 | 14 | 0.02 |
| AML_17 | BCORL1 | X | 129148573 | 129148573 | C | T | c.C1825T | p.R609X | nonsense | 183 | 389 | 0.68 |
| AML_22 | BCORL1 | X | 129155008 | 129155008 | - | G | c.3491dupG | p.R1164fs | frameshift-ins | 630 | 227 | 0.26 |
| AML_24 | BCORL1 | X | 129159171 | 129159171 | C | T | c.C3895T | p.R1299X | nonsense | 73 | 443 | 0.86 |
| AML_28 | BCORL1 | X | 129159270 | 129159270 | C | T | c.C3994T | p.R1332X | nonsense | 444 | 393 | 0.47 |
| AML_29 | BCORL1 | X | 129159270 | 129159270 | C | T | c.C3994T | p.R1332X | nonsense | 822 | 71 | 0.08 |
| AML_30 | BCORL1 | X | 129149098 | 129149098 | C | T | c.C2350T | p.R784X | nonsense | 442 | 67 | 0.13 |
| AML_30 | BCORL1 | X | 129159150 | 129159150 | C | T | c.C3874T | p.R1292X | nonsense | 447 | 23 | 0.05 |
| AML_33 | BCORL1 | X | 129159072 | 129159072 | C | T | c.C3796T | p.R1266X | nonsense | 541 | 13 | 0.02 |
| AML_34 | BCORL1 | X | 129171479 | 129171479 | - | GCTCATAGT<br>TCTTT | c.4443_4444insGCTCATA<br>GTTCTTT | p.N1481fs | frameshift-ins | 461 | 47 | 0.09 |
| AML_03 | KDM2B | 12 | 121879036 | 121879036 | C | T | c.G3078A | p.W1026X | nonsense | 279 | 229 | 0.45 |
| AML_25 | PCGF1 | 2 | 74732752 | 74732752 | C | A | c.G578T | p.R193L | missense | 52 | 415 | 0.89 |

**Supplemental Table S2.** Related to Figure 1D. Variant allele frequencies (VAFs) of variants detected in 4 male patients from two timepoints. nd: not determined.

|  | Variants | Variant allele frequencies (VAFs) |  |
| --- | --- | --- | --- |
|  |  | MDS (d0) | AML (d197) |
| Patient 1 | BCOR_P744fs* | 0.15 | 0.40 |
|  | BCORL1_I1217fs* | 0.11 | 0.48 |
|  | ETV6_R359L | 0.05 | 0.27 |
|  | PTPN11_E76Q | nd | 0.10 |
|  | Variants | Variant allele frequencies (VAFs) |  |
|  |  | MDS-EB2 (d0) | AML (d230) |
| Patient 2 | PHF6_Y301* | 0.81 | 0.82 |
|  | RUNX1_V164F | 0.44 | 0.43 |
|  | BCOR_Q1322* | 0.76 | 0.81 |
|  | BCORL1_Q1546* | 0.64 | 0.79 |
|  | RUNX1_S314fs* | 0.30 | 0.41 |
|  | NRAS_G13D | 0.31 | 0.44 |
|  | BCOR_K395* | 0.11 | nd |
|  | RUNX1_R166G | 0.06 | nd |
|  | Variants | Variant allele frequencies (VAFs) |  |
|  |  | MDS-EB2 (d0) | AML (d196) |
| Patient 3 | DNMT3A_R882H | 0.13 | 0.37 |
|  | U2AF1_S34F | 0.11 | 0.35 |
|  | BCOR_R1547* | 0.28 | 0.71 |
|  | RUNX1_S410* | 0.14 | nd |
|  | BCORL1_S1679fs* | nd | 0.61 |
|  | RUNX1_Y189fs* | nd | 0.36 |
|  | CSF3R_Q768* | nd | 0.40 |
|  | CSF3R_Q768* | 0.13 | 0.37 |
|  | Variants | Variant allele frequencies (VAFs) |  |
|  |  | MDS-EB2 (d0) | AML (d145) |
| Patient 4 | DNMT3A_R882C | 0.27 | 0.32 |
|  | BCOR_T819fs* | 0.60 | 0.65 |
|  | BCORL1_R1048* | 0.34 | 0.73 |
|  | NRAS_G12V | 0.08 | 0.32 |
|  | CUX1_E124fs* | 0.17 | nd |

**Supplemental Table S3.**
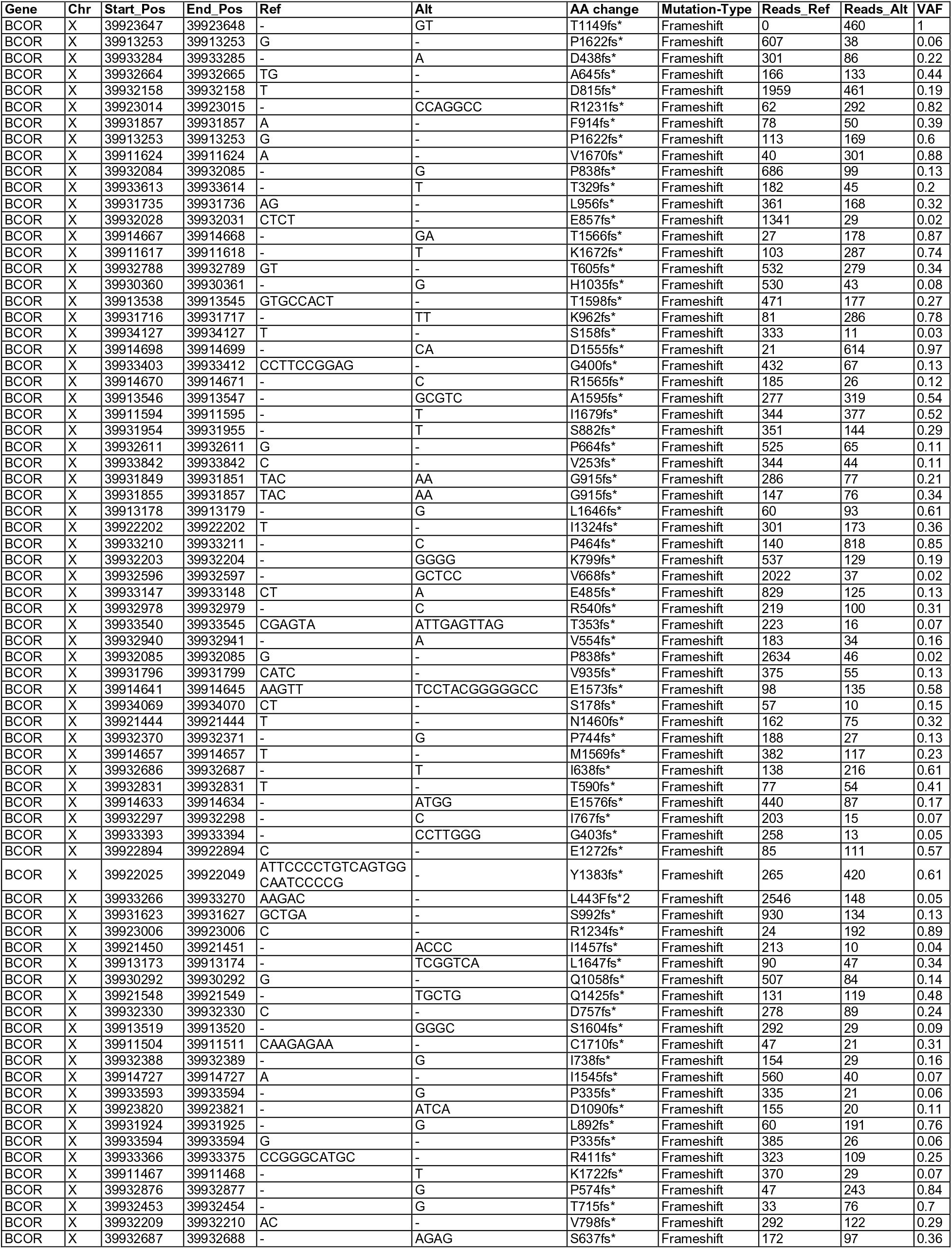
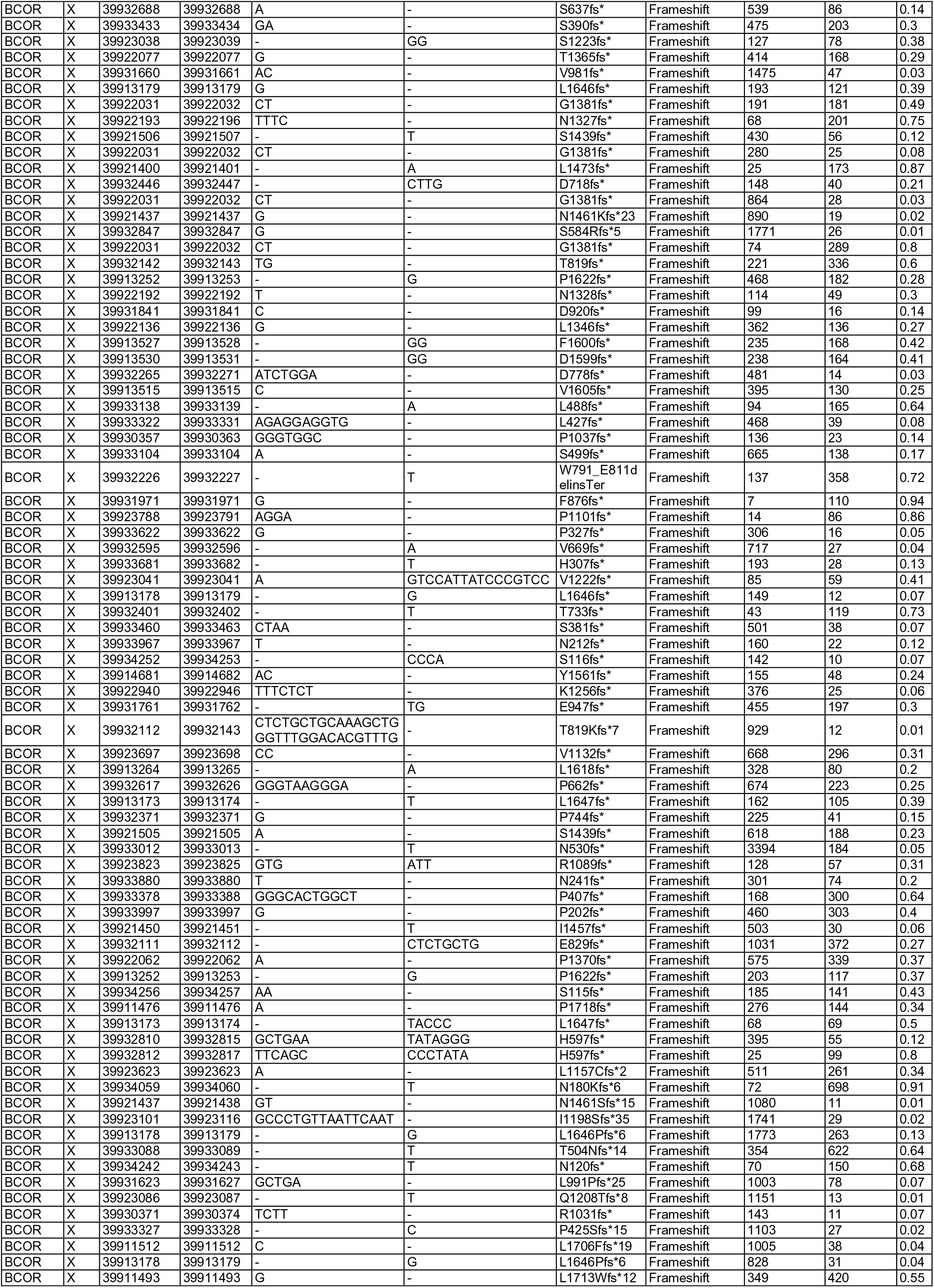
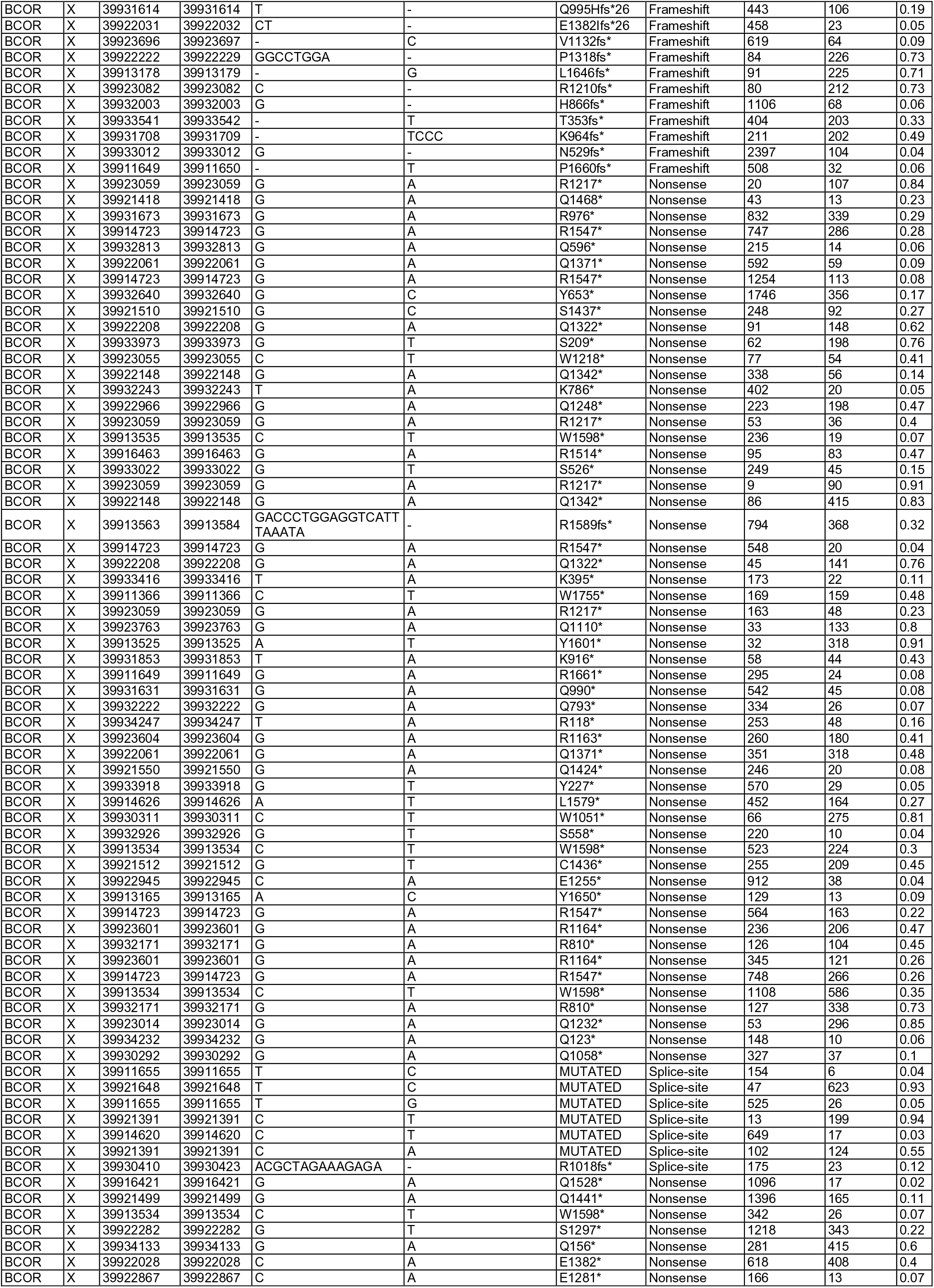
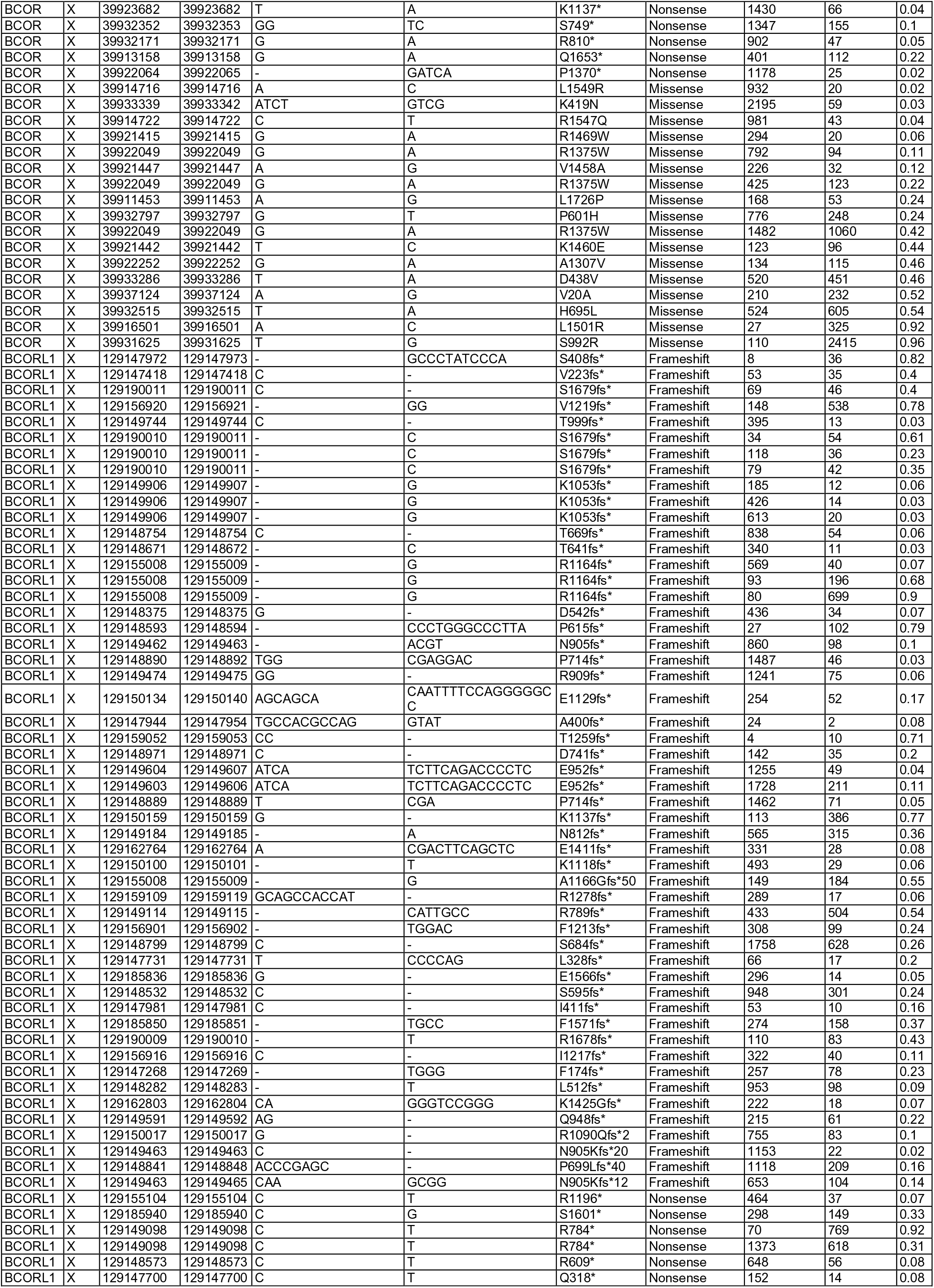
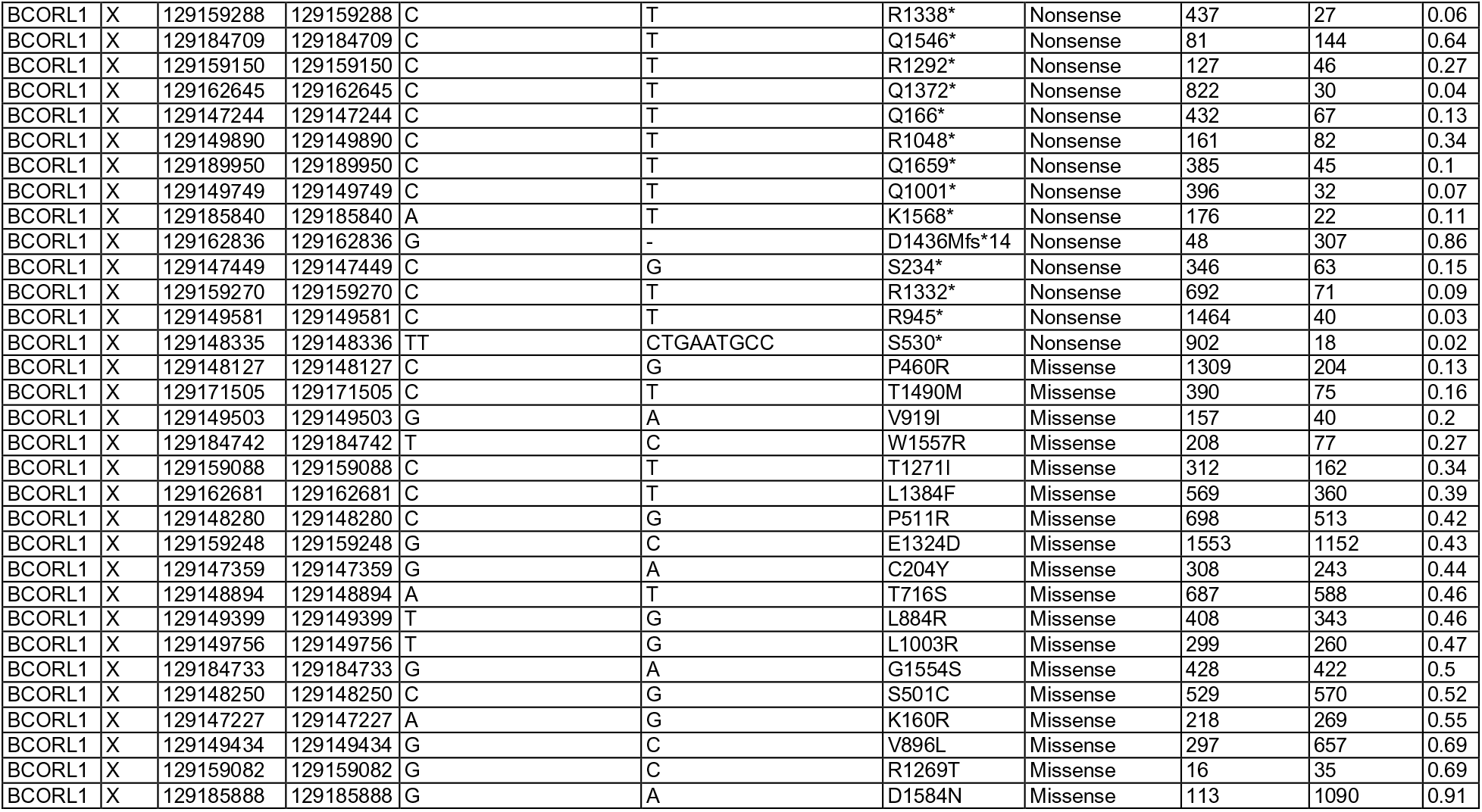
List of *BCOR* (n=256) and *BCORL1* (n=90) variants.

